# Molecular liver fingerprint reflects the seasonal physiology of the grey mouse lemur (*Microcebus murinus*) during winter

**DOI:** 10.1101/2021.12.22.473832

**Authors:** Blandine Chazarin, Margaux Benhaim-Delarbre, Charlotte Brun, Aude Anzeraey, Fabrice Bertile, Jérémy Terrien

## Abstract

Grey mouse lemurs (*Microcebus murinus*) are a primate species exhibiting strong physiological seasonality in response to environmental energetic constraint. They notably store large amounts of lipids during early winter (EW), which are thereafter mobilized during late winter (LW), when food availability is low. In addition, they develop glucose intolerance in LW only. To decipher how the hepatic mechanisms may support such metabolic flexibility, we analyzed the liver proteome of adult captive male mouse lemurs, which seasonal regulations of metabolism and reproduction are comparable to their wild counterparts, during the phases of either constitution or use of fat reserves. We highlight profound changes that reflect fat accretion in EW at the whole-body level, however, without triggering an ectopic storage of fat in the liver. Moreover, molecular regulations would be in line with the lowering of liver glucose utilization in LW, and thus with reduced tolerance to glucose. However, no major regulation was seen in insulin signaling/resistance pathways, which suggests that glucose intolerance does not reach a pathological stage. Finally, fat mobilization in LW appeared possibly linked to reactivation of the reproductive system and enhanced liver detoxification may reflect an anticipation to return to summer levels of food intake. Altogether, these results show that the physiology of mouse lemurs during winter relies on solid molecular foundations in liver processes to adapt fuel partitioning while avoiding reaching a pathological state despite large lipid fluxes. This work emphasizes how the mouse lemur is of primary interest for identifying molecular mechanisms relevant to biomedical field.

## Introduction

The measurement of time is a vital necessity for all living systems to be synchronized with the astronomical cycles of their environment. Each organism, from unicellular organisms to humans, harbors an endogenous biological clock allowing it to anticipate and adapt its biological functions to daily and seasonal cycles [1]. In nature, organisms use clues from their environment as indicators of time. The variations in sunlight constitute a precise and reproducible temporal signal which provides information on both the time of day (light intensity, light/dark cycles) and the day of year (day-length also called photoperiod). Also, this physical parameter has been selected during evolution as the most robust external indicator of time, unlike ambient temperature and food availability, which certainly influence biological rhythms, but to a lesser extent than photoperiod [2]. Mechanistically, the integration of photoperiod occurs through the secretion of melatonin by the pineal gland [2], and results in the generation of circannual rhythms. This feature is particularly critical in highly seasonal environments, characterized by drastic changes in abiotic (ambient temperature, rainfall) and biotic (energy availability) factors throughout the year.

Resource/energy availability fluctuates over the year, alternating between periods of food abundance and scarcity. To cope with these fluctuations, seasonal organisms alternate between active and inactive reproductive states during the favorable and unfavorable periods, respectively [3]. The reproductive seasonal phenotype of alternation between periods of gonadal recrudescence and regression, to be compared to yearly puberty, is accompanied by profound changes in metabolic status, generally reinforced by the necessity to respond to harsh environmental conditions during winter. Seasonal reproduction is indeed often characterized by extreme winter metabolic phenotypes, including, e.g. migration, hibernation and daily torpor. Physiological remodeling underlining such changing phenotypes includes changes in lipid metabolism, cell membrane function, neural and cardiovascular functions, anorexia- and hypoxia-related responses, skeletal muscle function, immune function and non-shivering thermogenesis [4,5].

Non-model organisms or exotic species could be highly valuable as their natural life-history traits have evolved into behavioral and physiological adjustments to repeatedly survive periods of food scarcity and abundance. Such an evolution ended with the expression of adaptive phenotypes revealing striking features that may be of extreme importance in a clinical context [6–10]. More specifically, seasonal species undergo massive fattening to anticipate periods of partial or complete energy depletion in the field. This ability to store massive amounts of fat in anticipation to phases of poor food availability seems associated with protective mechanisms against adverse effects usually associated with alterations of glucose/insulin signaling [11,12]. Although energy homeostasis relies on a complex set of mechanisms, involving multiple neuroendocrine pathways [13] and centered on hypothalamic control [14,15], it is also quite clear that peripheral tissues, including liver, pancreas, muscle and adipose tissue, play major roles in the seasonal regulation of energy homeostasis, especially at the level of nutrient partitioning [16]. However, the molecular bases of seasonal adaptation are not always fully understood, e.g., in the liver, a key tissue to metabolic flexibility [17], i.e. the capacity to switch from carbs to lipid oxidation.

The grey mouse lemur (*Microcebus murinus*) is a nocturnal lemuriform primate (Cheirogaleidae), phylogenetically close to humans and showing robust seasonal modifications in energy balance related to its endemicity to Madagascar. Relative to its body size (body mass varying between ~50 and ~90g in wild conditions), this species has a long lifespan (half-life around 5.7 years; mean lifespan ~8-10 years in captivity [18]), which makes it a good model for ageing studies [19]. In addition, mouse lemurs undergo profound seasonal remodeling synchronized on photoperiod [20] and following annual variations in environmental conditions, including rainfalls, ambient temperature and consequently food availability [21,22]. Energy homeostasis in the mouse lemur is characterized by spontaneous transitions from high to low body mass levels along seasons, resulting from an exceptional capacity to adjust energy metabolism to food scarcity during winter or high food availability during summer [23]. In particular, the extensive use of daily torpor in early winter (EW) allows the mouse lemur to build up fat reserves which are later mobilized for reproduction, therefore conferring to winter fattening a strong adaptive value. More specifically, the condition for reproductive success in the male is centered on access to females, which constitutes its major energy constraint. Therefore, male mouse lemurs must reactivate their reproductive system upstream of the female receptivity period, therefore anticipating the transition from winter to summer. Male mouse lemurs’ reproductive investment indeed relies on spermatogenesis and mating with competition, leading to massive energy investment in late winter (LW), when energy is not available in the environment.

The seasonal phenotype in mouse lemurs can be mimicked in captivity with the only manipulation of photoperiod (to reproduce the seasonal transitions) and is completely comparable to what is observed in wild individuals, although the environmental constraint is drastically reduced. Indeed, captive male mouse lemurs acclimated to winter-like photoperiod (10 hours / day) during 6 months and maintained in constant housing conditions (high, stable ambient temperature, food availability) exhibit deep changes in body mass explained by variations in metabolic rates, thus resulting in a biphasic phenotype, with a phase of massive body mass gain in EW, followed by the spontaneous arrest of fattening in LW [11]. Strikingly, mouse lemurs are protected against glucose intolerance during fattening, but not during the second half of winter when they lose fat and exhibit increased fasting insulin levels. Most importantly, these natural transitions are fully reversible, and never reach a pathological stage, as reflected by the apparent absence of an inflammatory response [11]. With regard to classical models of obesity, these results emphasize a paradox as insulin sensitivity seems to be preserved during the fattening phase. However, a similar paradox was described in hibernating bears that preserve insulin sensitivity during the fattening period through altered adipose PTEN/AKT signaling [24]. Finally, recent work has shown that the relative expression of phospho-IRS-1 was enhanced during torpor in the mouse lemur in muscle, while phospho-insulin receptor level was decreased in white adipose, thus suggesting an inhibition of insulin/IGF-1 signaling during torpor in these tissues [25]. These results altogether show that mouse lemurs undergo profound physiological changes underlying their ability to spontaneously reverse the fattening process. We investigated which liver pathways are activated/repressed at the protein level in the fattening (EW) vs slimming (LW) phenotype of mouse lemurs by comparing phases of gain and loss of body mass, using a mass spectrometry-based approach. We expected major regulations of the glucose/insulin and lipid pathways, according to the physiological phenotype described in the paper by Terrien et al. [11].

## Materials and methods

### Animals and ethical statement

Adult male grey mouse lemurs (mean age ± SEM: 3.7 ± 0.6 yo), all born in the captive population of Brunoy (MNHN, France, license approval N° E91.114.1), were randomly chosen used and were in good health at the start of the experiment. General conditions of captivity were maintained constant: Ta (24 - 26 °C), relative humidity (55%). Food (including fresh fruits and a lab-made mixture) and water were available ad libitum. Diet (1.31 kcal/g) was composed of cereals, gingerbread, condensed milk, yogurt, banana and egg and resulted in a macronutrient balance of ~60% carbohydrates, ~20% proteins and ~20% lipids. Diet fatty-acids were distributed as previously described [26]. In captivity, seasonal variations of physiological functions can be entrained by alternating 6-month periods of summer-like long photoperiod (14 h of light / day) and winter-like short photoperiod (10 h of light / day) under artificial light. The present study focused on winter season such as adult (mean age ± SEM: 4.1 ± 0.1 years, range: 3.3–4.3) male mouse lemurs were maintained in constant photoperiod conditions (10 h of light / day). General conditions of captivity were applied, and animals were maintained in social groups. All experiments were carried out in accordance with the European Communities Council Directive (86/609/EEC) and all experimental procedures were evaluated by an independent ethical council and approved for ethical contentment by the French government (authorizations n°03210.02, n°APAFIS#3004-2015111015031850 and n°APAFIS#3697-2016012111304236).

### Glycaemia, circulating lipids and metabolic parameters

Fasting glycemia (mg*dL^−1^) using a hand-held blood and circulating lipids (total cholesterol CHOL, high-density lipoprotein HDL and triglycerides TG, mg*dL^−1^), using a hand-held blood OneTouch® Vita glucometer (LifeScan, France) and Mission® Cholesterol Meter (ACON Laboratories, Inc., USA), respectively, were measured after blood puncture from the saphenous vein during the resting phase of the animals (Early Winter EW, N=9; Late Winter LW, N=10) and before food became available. Low-density lipoprotein levels (LDL, mg*dL^−1^) and CHOL/HDL ratios were calculated. In addition, metabolic data were reanalyzed from already published [details on the methods in 11] and additional unpublished work to extract data from individuals exhibiting the most extreme phenotypes between EW (fattening: highest body mass gain; N=11, BMC range=+19-39%, median=22%) and LW (leaning: highest body mass loss; N=9, BMC range=−10-28%, median=−14%) winter. Data [details of the methods in 11] on food efficiency (kcal/g), metabolic performance (nocturnal mean values of VO2 in ml/kg/h, heat production in kcal/hr and respiratory exchange ratio RER) and glucose homeostasis (area under the curve AUC in arbitrary units during glucose tolerance test and basal insulinemia in ng/mL) were all put in perspective of observations made in the proteomic analysis.

### Body mass and liver biopsies

Five animals were randomly chosen for each critical period of winter season (N=10 liver samples total): during the first half of the winter (week 6 of 26, 20% of winter period undergone, EW) and the second half of winter (weeks 18 of 26, 66% of winter period undergone, LW). Body mass (BM) was recorded at the time of liver biopsy. In addition, animals are continuously monitored for body mass in the captive population and data are implemented in a database. These data were used to calculate the difference between the body mass (BMC; expressed in %) recorded at the onset (for animals sampled in EW) or the offset (for animals sampled in LW) of the 6-month winter photoperiod and the maximum body mass (usually reached in mild winter).

Liver biopsies (5-10mg samples) were sampled at the apex of liver in mouse lemurs under general anesthesia (induction with Diazepam 0.05 mg/kg, Valium®, Roche; Ketamine 0.2 mg/kg, Imalgen 1000, Boehringer Ingelheim; maintenance with isoflurane 1-2 %, Iso-Vet, Piramal Healthcare) and per-operating analgesia (before surgery: buprenorphine 0.05 mg/kg IM 30 minutes, local cutaneous injection of lidocaine around the abdominal aperture; after surgery: renewal of buprenorphine 4 hours later, then meloxicam *per os* for 2 days). Monitoring was maintained over two weeks to ensure that the animal healed and recovered properly.

### Liver gene expression

Hepatic gene expression was investigated using qPCR arrays designed for human (RT^2^ Profiler PCR arrays – ‘Human Fatty Liver’, Qiagen France SAS). Then, mRNA was extracted using trizol reagent and following the instructions of the manufacturer. RNA concentration was determined using Qubit fluorometric quantitation (Thermo Fisher Scientific Inc) and contamination of RNA was assessed using 2100 Bioanalyzer instrument (Agilent Technologies, Santa Clara, CA, USA). At this step, one sample was determined as contaminated (sample from the second half of winter) and therefore excluded from the experiment. The reverse transcription and gene expression analysis were then processed following the instructions of the manufacturer. A pre-analysis using the Qiagen online PCR Array Data Analysis Software determined that 1 of our samples was not proper for gene expression analysis which meant that we had 4 liver samples left for both the first and second half of winter. Variations between early and LW have already been published [11] and will not be discussed here. Instead, gene expression data will be used to test the existence or not of any correlation between proteomic and transcriptomic results as both methods have been run on the same liver samples.

### Liver proteome analysis

Unless otherwise specified, all reagents and chemicals were purchased from Sigma Diagnostics (St. Louis, MO, USA). Frozen liver samples were weighed and grinded using a ball mill (2 × 30 s @ 25 Hz; MM400, Retsch) under liquid nitrogen and proteins were extracted using 10 volumes (i.e. 10 μL/mg of tissue) of extraction buffer (8 M urea, 2 M thiourea, 2% CHAPS, 2% DTT, 30 mM Tris pH 8.8, protease inhibitors). After sonication on ice (3 × 10 sec., 135 Watts), followed by centrifugation (10 min, 2000g, 4°C) to remove possible cell debris, proteins were acetone-precipitated (6 volumes of cold acetone) overnight at −20°C. Protein pellets were obtained by centrifugation (10 min, 13500g, 4°C) before being washed using cold acetone and re-solubilised in Laemmli buffer (10 mM Tris pH 6.8, 1mM EDTA, 5% β-mercaptoethanol, 5% SDS, 10% Glycerol). Protein concentrations were determined using the RC-DC Protein Assay (BioRad, Hercules, CA, USA), and similarity of protein electrophoretic profiles between all samples was then confirmed using 12% SDS-PAGE acrylamide gels (20 μg loaded) using colloidal Coomassie blue (Fluka, Buchs, Switzerland) staining. At this stage, a reference sample comprising equal amounts of all protein extracts was made, to be repeatedly analysed during the whole duration of nanoLC-MS/MS experiments for allowing QC-related measurements.

Fifty μg of each sample were mixed with 4x SDS sample buffer (1x: 10 mM Tris pH 6.8, 1 mM EDTA, 5 % β-mercaptoethanol, 5 % SDS and 10 % glycerol), incubated at 95 °C for 5 minutes and proteins were separated by electrophoresis using SDS-PAGE gels composed of a stacking (5 % polyacrylamide) and a running (12 % polyacrylamide) layer. Electrophoresis was stopped after protein migration in the running gel had reached 1.2 cm, and protein staining was performed using colloidal Coomassie Blue. Six gel bands (2 mm each) were excised, and proteins were in-gel digested with trypsin (Promega, Madison, WI, USA; ratio 1:100) at 37°C overnight after de-staining, reduction (10 mM DTT), alkylation (55 mM iodoacetamide), and dehydration using a MassPrep station (Micromass, Waters, Milford, MA, USA). Tryptic peptides were extracted by incubation on an orbital shaker (450 rpm) for one hour using 60% acetonitrile, 0.1% formic acid in water and then for 30 minutes using 100% acetonitrile. A set of reference peptides (iRT kit; Biognosys AG, Schlieren, Switzerland) was finally added to all samples. Solvent was then eliminated using a vacuum centrifuge (SpeedVac, Savant, Thermoscientific, Waltham, MA, USA), and peptides were re-suspended in 30 μL of 1% acetonitrile, 0.1% formic acid in water.

Tryptic peptides were analysed on a nanoUPLC system (nanoAcquityUPLC, Waters, USA) coupled with a quadrupole-Orbitrap hybrid mass spectrometer (Q-Exactive Plus, Thermo Scientific, San Jose, CA). The six peptide extracts from an individual sample were injected in a row, and samples from the two animal groups were injected alternatively. To minimize carry-over, a column wash program (50% acetonitrile during 20 min) and two solvent blank injections were included in between each biological replicate. Peptide extracts from the reference sample were injected three times over the course of nanoLC-MS/MS analyses.

Briefly, 1.5 μl of each sample were concentrated/desalted on a trap column (Symmetry C18, 180 μm x 20 mm, 5 μm; Waters) using 99 % of solvent A (0.1 % formic acid in water) and 1 % of solvent B (0.1 % formic acid in acetonitrile) at a flow rate of 5 μl/min for 3 minutes. Afterwards, peptides were eluted from the separation column (BEH130 C18, 75 μm x 250 mm, 1.7 μm; Waters) maintained at 60°C using a 65 min gradient from 1-35% of B at a flow rate of 450 nL/min.

The Q-Exactive Plus was operated in positive ion mode with source temperature set to 250°C and spray voltage to 1.8 kV. Spectra were acquired through automatic switching between full MS and MS/MS scans, in data dependent acquisition mode. Full scan MS spectra (300-1800 m/z) were acquired at a resolution of 70,000 (m/z 200) with an automatic gain control (AGC) value set to 3 × 10^6^ ions, a maximum injection time set to 50 ms, and the lock-mass option being enabled (polysiloxane, 445.12002 m/z). Up to 10 most intense multi-charged precursors per full MS scan were isolated using a 2 m/z window and fragmented using higher energy collisional dissociation (HCD, normalised collision energy of 27 eV). Dynamic exclusion of already fragmented precursors was set to 60 seconds. MS/MS spectra were acquired at a resolution of 17,000 (m/z 200) with an AGC value set to 1 × 10^5^, a maximum injection time set to 100 ms, and the peptide match selection option was turned on. The system was fully controlled by Xcalibur software (v3.0.63; Thermo Fisher Scientific). Peak intensities and retention times of reference peptides were monitored in a daily fashion.

MS raw data were processed using MaxQuant (v1.5.3.30). Peak lists were created using default parameters and searched using Andromeda search engine implemented in MaxQuant against a protein database derived from the latest annotation of the *Microcebus murinus* (TaxID 30608) genome in Refseq (Refseq Assembly accession GCF_000165445.2; Assembly Name Mmur_3.0). Only the longest proteins per gene (coding DNA sequences) were retained, and after elimination of any redundancy, the database contained 39,712 protein sequences to which sequences of common contaminants were added (247 entries; contaminants.fasta included in MaxQuant). The first search was performed using a precursor mass tolerance of 20 ppm, and 4.5 ppm for the main search after recalibration. Fragment ion mass tolerance was set to 20 ppm. The second peptide research option was enabled. Carbamidomethylation of cysteine residues was considered as a fixed modification and oxidation of methionine residues and acetylation of protein N-termini as variable modifications during the search. A maximum number of two missed cleavages and a false discovery rate (FDR) of 1% for both peptide spectrum matches (minimum length of seven amino acids) and proteins were accepted during identification. Regarding quantification, data normalisation and protein abundance estimation were performed using the MaxLFQ (label-free quantification) option implemented in MaxQuant [27] using a “minimal ratio count” of one. “Match between runs” was enabled using a two-minute time window after retention time alignment. For quantification, we considered unmodified peptides and, if detected, also their modified counterpart (acetylation of protein N-termini and oxidation of methionine residues), while shared peptides were excluded. All other MaxQuant parameters were set as default. Proteins identified with a single peptide were not considered, and eventual contaminants and reversed proteins were removed from the dataset. Only proteins with at least four of five values per group as well as the ones “absent” (i.e. zero valid values) in samples from a given group were kept for quantification. The mass spectrometry proteomics data have been deposited to the ProteomeXchange Consortium via the PRIDE partner repository with the dataset identifier PXD030198.

To annotate identified *Microcebus murinus* proteins, we searched known homologous proteins in the human Swissprot database (accessed in April 2019). This was done by using BLAST searches (FASTA program v36; downloaded from http://fasta.bioch.virginia.edu/fasta_www2/fasta_down.shtml), and only the top BLAST hit per protein was retained. To validate this procedure, we automatically extracted orthology annotations of *Microcebus murinus* proteins and of their human homologues from the OrthoDB resource [28]. The relevance of the match between *Microcebus murinus* proteins and their human homologues was then checked manually.

### Proteomic bioinformatic analysis

To identify functionally relevant biological processes differently regulated in EW and LW animals, enrichment and functional annotation analyses of differential proteomics data were performed using two complementary algorithms. First, we used the desktop version of DAVID (Ease v2.1) and a version of Gene Ontology (GO) and Kyoto Encyclopedia of Genes and Genomes (Kegg) databases downloaded in July 2019. Stringent criteria for considering significantly enriched GO/Kegg terms included an Ease score lower than 0.001, a Benjamini p-value lower than 0.01, and a fold enrichment higher than 2. The liver functions altered between EW and LW were determined after enriched GO/Kegg terms had been grouped together into broad functional categories. Secondly, we used the interactome analysis and CAME (ID Conversion, Annotation, Membership, function Enrichment) workflow for annotation enrichment analysis applied in Metascape (https://metascape.org/). Default parameters were used. Using mainly Biogrid database, information about physical protein-protein interaction was first extracted and a protein interaction network was built. Molecular Complex Detection (MCODE) algorithm was then applied to this network to identify neighborhoods where proteins were densely connected. Ontology enrichment analysis, which was performed using the most popular annotation sources, e.g., GO and Kegg, but also WikiPathways, Reactome, Corum, and a series of other databases, was finally applied to each MCODE neighborhoods to assign them a biological function on the basis of the top-three (when available) functional enriched terms, when available. Networks were visualized using Cytoscape software v3.8.2.

### Western-blot analysis

The liver proteins previously solubilized (see above) in Laemmli buffer were used for Western blotting. Total protein concentration was determined using the Bio-Rad RC DC Protein Assay kit (Hercules, CA, USA). Protein electrophoresis was performed by loading 20 μg of each sample onto 4–20% Bio-Rad Mini-Protean TGX Stain-Free Precast Gels (Bio-Rad). Gel imaging was obtained after activation using the Bio-Rad ChemiDoc Touch Imaging System, then proteins were transferred to 0.2 μm nitrocellulose membranes (Bio-Rad) using the Trans-Blot Turbo Transfer System (Bio-Rad). Blot imaging allowed checking for the quality of the transfer. Membranes were blocked for 1 hour at room temperature with a solution of TBS-T (Tris 25 mM, NaCl 137 mM, KCl 2.68 mM, 5% Tween 20) containing 4% of BSA, and they were then incubated overnight at 4°C with primary antibodies. Primary antibodies were purchased from ThermoFisher Scientific (Waltham, MA, USA; Invitrogen manufacturer) for mitochondrial hydroxyacyl-coenzyme A dehydrogenase (HADH; PA5-31157) and citrate synthase (CS; PA5-221126) and from abcam (Paris, France) for uncoupling protein 3 (UCP3; ab10985). They were used at 1/2000 (HADH), 1/1000 (CS), and 1/500 (UCP3) dilution in the blocking solution. Membranes were then washed three times for 10 minutes in TBS-T, before being incubated for 1 hour at room temperature with a peroxidase-conjugated secondary antibody purchased from Santa Cruz Biotechnology (Santa Cruz, CA, USA), which was used at 1/5000 dilution in the blocking solution. Membranes were then washed three times for 10 minutes in TBS-T, before being incubated for 5 secondes in Luminata Classico Western HRP substrate (Merck Millipore, Molsheim, France) and then imaged for chemiluminescence using the ChemiDoc Touch Imaging System (Bio-Rad). Image analysis was done using the Bio-Rad Image Lab software v5.1. Signals were normalized to total proteins, as measured from stain-free gel images, and intensity values were expressed relative to those from EW animals, which were assigned an arbitrary value of one.

### Statistical analysis

Body mass and metabolic data are given as means ± standard error to the mean (SEM) and statistical significance was obtained using t-tests (P-values < 0.05). Statistical analysis of proteomics quantitative data was performed under R software environment v3.4.0 [29]. Normality of MaxLFQ values distribution and homoscedasticity were checked using Shapiro-Wilk and Bartlett tests (P-values > 0.01), respectively. Changes in protein abundances between animal groups were tested using Welch Student t-tests, significance being set to P-values < 0.05 (given the small number of animals, we sometimes extended our data exploration to a significance level set at p < 0.1). A principal component analysis (PCA) was computed with all quantified proteins using the ‘FactominR’ 2.4 package in R. Missing values were imputed using the ‘missMDA’ 1.18 package. A heatmap was generated based on differentially expressed proteins between EW and LW (p-value < 0.05) using the ‘heatmap.2’ function in the ‘gplots’ 3.1.1 package. Hierarchical clustering was applied within animals only, not proteins, to attest for the difference between EW and LW animals, based on their proteomic profiles.

## Results

### Same body mass but different dynamics

Mouse lemurs sampled during the first half of winter (Early Winter, EW) exhibited the same mean body mass than those sampled during the second half of winter (Late Winter, LW) (mean ± SEM: 104.0 ± 4.7g vs 101.4 ± 6.0g in EW and LW, respectively; t=0.3, p=0.74; Figure 1A). However, the analysis of body mass variations revealed striking differences between EW and LW. Indeed, animals sampled in EW (i.e. 6 weeks after winter onset) were actually undergoing massive fat accretion and had gained 55 ± 11% BM during the first half of winter (Figure 1B). In comparison, mouse lemurs sampled in LW had lost 37 ± 5% BM, therefore showing strong fat mobilization, during the second phase of winter (t=6.8, p<0.001; Figure 1B), thus confirming that animals in EW and LW were undergoing very distinct body mass dynamics. In this line, mouse lemurs from the LW group showed lower circulating levels of triglycerides (TG; −36%, p=0.001) but greater levels of total cholesterol (CHOL) and low-density lipoprotein (LDL) (+33%, p=0.001 and +51%, p=0.0001, respectively) and a tendency for a higher CHOL/HDL ratio (+32%, p=0.102) than EW animals, while glycaemia and high-density lipoprotein (HDL) remained unchanged (Figure 1C). These changes showed interesting correlations with BM variations (Figure 1D): animals showing the highest body mass loss were those with the lowest circulating levels of TG (r^2^=0.31, p=0.01) but the highest levels of CHOL (r^2^=0.21, p=0.05) and LDL (r^2^=0.32, p=0.02).

**Figure 1.**
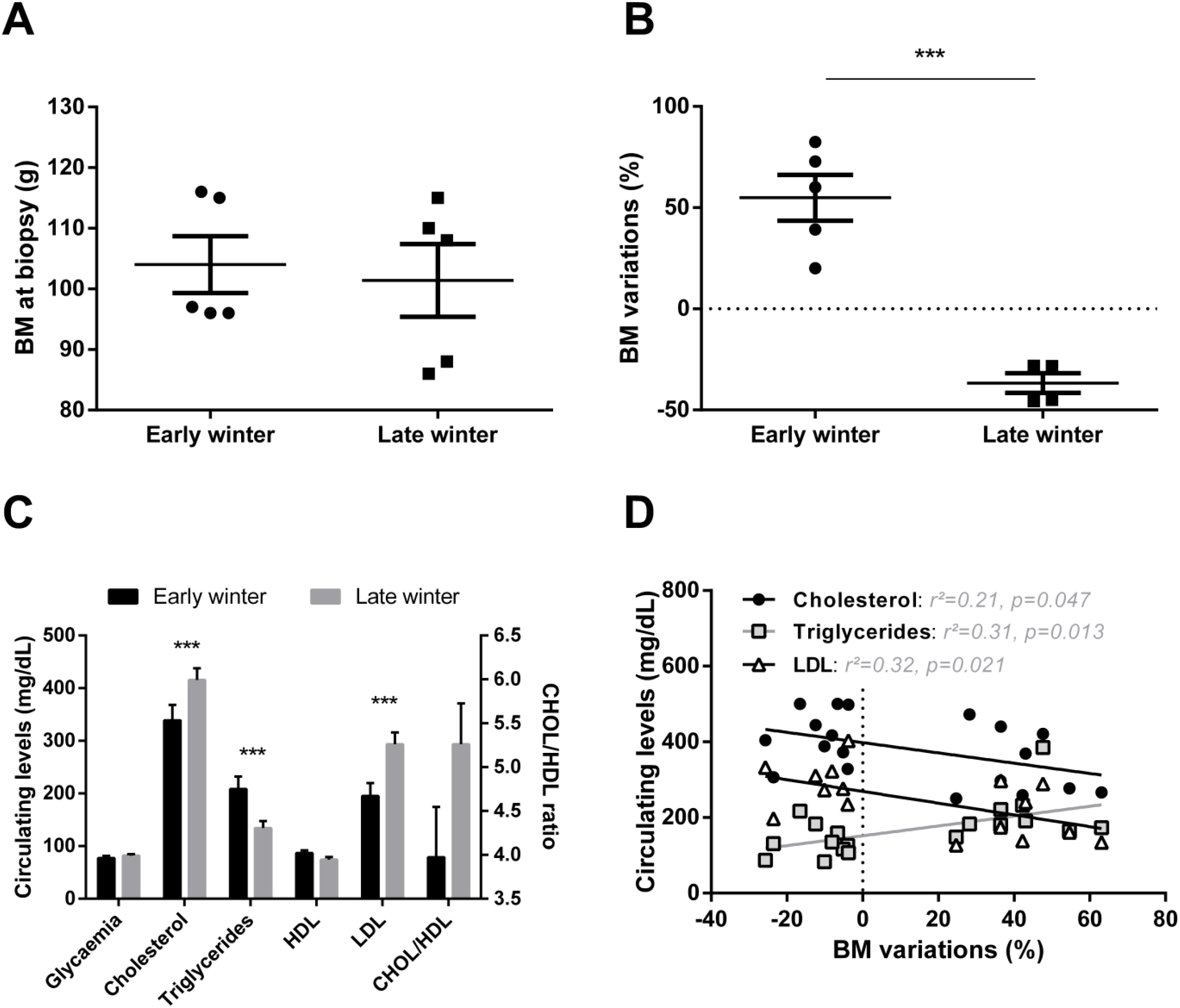
Body mass and plasma profiles in grey mouse lemurs during winter. Although mouse lemurs sampled in early (EW) and late (LW) winter (N=5/group) exhibited similar body masses (BM) at the time of liver biopsy (panel A), variations during the first vs second half of winter clearly showed BM gain in EW but BM loss during LW (panel B). Changes in BM dynamics were paralleled by changes in plasma metabolites and lipoprotein levels (panel C, data are presented as the means ± SEM), the relationship between these parameters being significant in some cases (panel D). *** indicate significance at p < 0.0001 between EW and LW.

### Overview of liver proteomics data

HPLC performances remained very good and stable throughout the whole experiment, as median coefficient of variation (CV) of retention times of all iRT peptides when considering all injections was 0.7%. The reproducibility of quantitative data was also very good, with low median CVs for the raw intensity of iRT peptides among all injections (13%), and for Maxquant-derived LFQ values of all quantified proteins within each of the two experimental groups (21.6%) and in the reference sample injected repeatedly during the course of MS-based analyses (9.5%).

Overall, the identification of 4,964 proteins was robust enough to comply with the stringent filtering criteria that were applied (see above), and 4522 of these proteins were retained for MS1 intensity-based label-free quantification after elimination of those exhibiting a single unique peptide (Table S1). Based on the expression of these 4522 proteins, very good discrimination could be observed between the two biological groups using a PCA analysis (Figure 2A). Indeed, animals from the EW and LW groups discriminated significantly around dimension 2 (r^2^=0.42, p=0.04), which was mostly explained by the regulations of 759 proteins, amongst which GAPDH and LDHA (involved in carbohydrate metabolism), MAN1B1 (involved in protein metabolism) and NSDHL (involved in lipid metabolism) were included in the 30 most contributory proteins (Figure 2A, Table S2).

**Figure 2.**
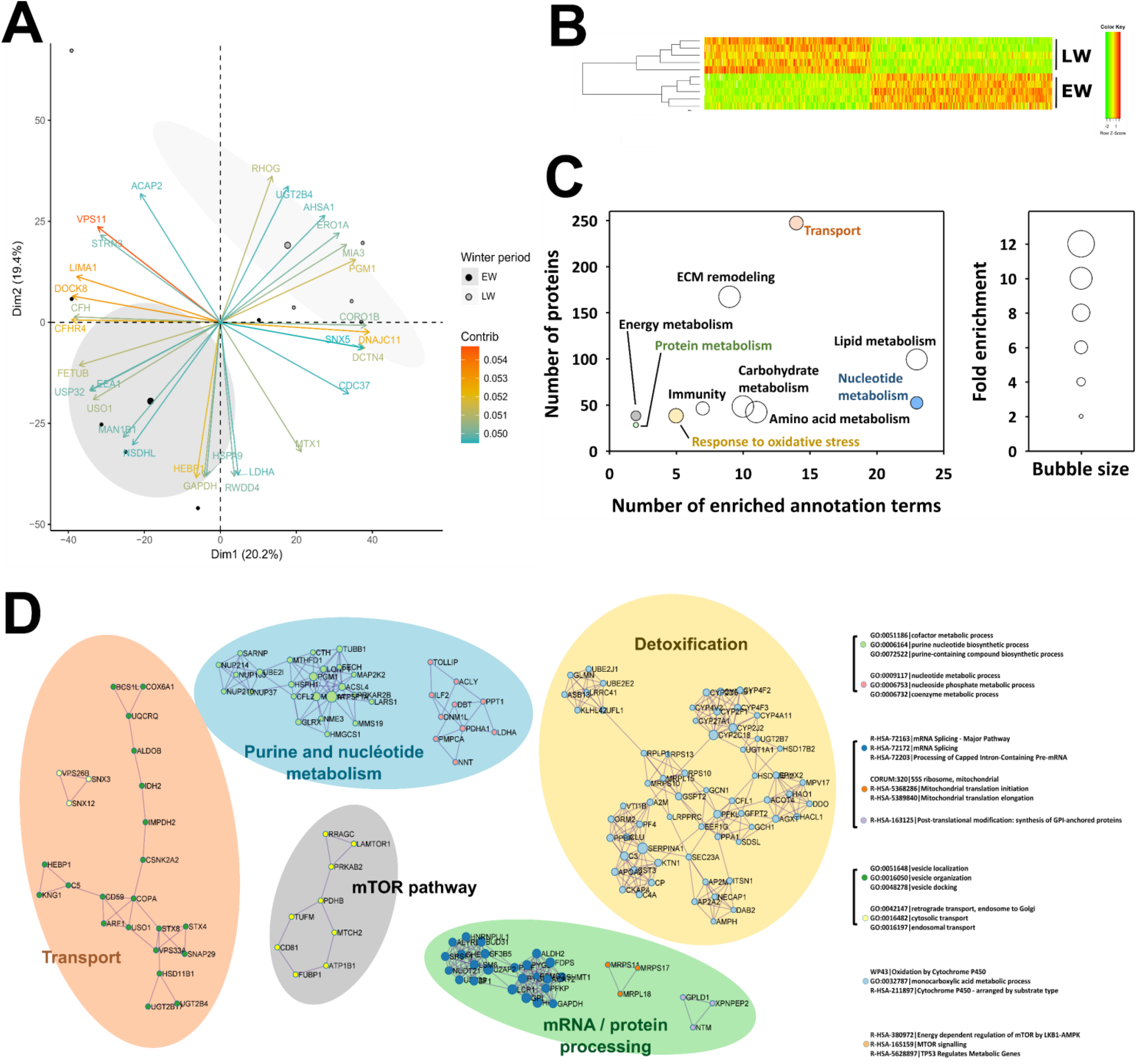
Overview of liver proteomics data. Principal component analysis performed from expression levels of all 4522 proteins analyzed using mass spectrometry (panel A; the 30 most contributing proteins are represented) and hierarchical clustering performed from expression levels of all 538 proteins found differentially expressed between early (EW) and late (LW) winter (panel B) allowed these two conditions to be well discriminated. Functional annotation analysis performed from the list of the 538 differential proteins revealed enriched ontology terms, which allowed determination of broad functions significantly affected between EW and LW. Complementary “functional information” was obtained using the algorithm from Ease/DAVID (panel C) and that from Metascape (panel D). Colors in panels C and D indicate overlapping information. Detailed protein abundances and fold changes are given in Table S1. ECM: extracellular matrix.

The abundance of 534 proteins was significantly changed between EW and LW, 288 of them being higher in the LW versus EW period, while the reverse was observed for the 246 remaining proteins. Three additional proteins (NCF1, HSD11B2, TMX2) were not detected in EW animals and one (CYP2C18) in LW. Also, PCA analysis (data not shown, PC1 at 59%) and hierarchical clustering (figure 2B) showed a distinct discrimination of the two biological groups confirming the low variability between biological replicates. Two different algorithms for functional annotation enrichment analysis from all differentially abundant proteins (including proteins not detected in a given phase) brought similar and complementary information. Using the EASE/DAVID algorithm, a total number of 106 GO terms/Kegg maps were found significantly enriched, which allowed highlighting ten processes, including lipid/carbohydrate/amino-acid metabolisms and the remodeling of extracellular matrix (ECM) as the most enriched (Figure 2C). Enrichment in annotation terms was also observed, to a lesser extent, for other functions like energy, nucleotide and protein metabolisms, transport, response to oxidative stress and immune response. Using Metascape/Cytoscape, 25 annotation terms were found significantly enriched, corresponding to differential proteins distributed in nine interaction networks that define five functional categories, including detoxification, nucleotide metabolism, transport, mRNA processing and protein metabolism, and finally the mTOR pathway (Figure 2D).

### Specific analysis of significantly enriched pathways

#### Protein metabolism

A high number of liver proteins involved in protein metabolism were detected. Comparing their abundance in LW vs EW, a p-value lower than 0.05 was calculated for sixty of them. Concerning protein synthesis, higher abundance of 6 proteins (SF1, SUPT4H1, SUPT5H, TCEA1, U2AF2, and USP39; +20 to +93%, median = +27%) in LW may support an overall higher transcriptional activity at this stage, while the reverse could be interpreted for lower abundance of 2 proteins (LSM6 and SF3B5; −4% and −13%, respectively). Protein translational process also appeared regulated during winter, with 9 proteins more abundant (GCN1, MRPL15, MRPL18, MRPS10, MRPS11, MRPS17, RPLP1, RPS10, and RPS13; +9 to +62%, median = +28%) and 12 other ones less abundant (CARS, CSNK2A2, EEF1G, EIF5, GSPT2, GATB, LARS, MTG1, OSGEP, RPS6KA1, TUFM, and UPF1; −12 to −45%, median = −20%) in LW versus EW. Finally, adjustments in the process of protein folding seem to be needed depending on winter timing, with the abundance of 1 protein being higher (DNAJB11, +44%), and that of 3 other ones being lower (MAN1B1, NUDCD2, and OS9; −18 to −32%, median = −25%) in LW versus EW animals.

Concerning protein catabolism, numerous changes were recorded for the ubiquitin proteasome system, with 8 proteins higher expressed (ATG3, GLMN, MARCH5, NUB1, PSMB10, PSME3, UBE2E2, UBE2J1, +29 to +90%; median = +40%) and 11 other ones lower expressed (ASB13, CUL4, KLHL42, LRRC41, PSMD2, PSMG2, PSMG3, PSMG4, UCHL5, UFL1, and USP32; −12 to −60%, median = −27%) in LW versus EW lemurs. With three proteins higher expressed (BNIP3, RMC1, and SNAP29; +43 to +61%, median = +60%), and only one lower expressed (NRBF2; −30%) in LW, liver autophagy may appear enhanced at this stage. Finally, expression levels of other liver intracellular proteases were found higher (DNPEP; +28%) and lower (NRDC, XPNPEP2, and XPNPEP3; −23 to −36%, median = −32%) in LW compared to EW. Changes in the abundance of proteins controlling protein metabolism were reinforced by the observation of 46 additional regulated proteins when significance level was set between 0.05 and 0.1 (see Table S1 for details). Among them, upregulated proteins in LW vs EW included 3 proteins involved in transcription (ELP2, RTRAF, and SUPT6H; +5 to +21%, median = +19%), 12 in translation (ATE1, BRIX1, EEF1E1, EIF1B, MRPL17, MRPL19, MRTO4, NOP58, PARS2, RARS2, RPS17, and WARS; +12 to +69%, median = +29%), 1 in protein folding (AUP1, +13%), 7 in the ubiquitin proteasome system (DTX3L, OTULIN, PSMA1, PSMB9, UBE2G2, UBE3C, USP19; +15 to +128%, median = +38%), 1 in autophagy (SQSTM1, +62%). Downregulated proteins in LW vs EW grouped together one protein involved in transcription (SNU13, −11%), 9 in translation (AARSD1, EIF3A, EIF3J, MRPL2, MRPS21, RPL34, RPS27L, RPS28, and SBDS; −5 to −47%, median = −20%), and 12 in the ubiquitin proteasome system (BRCC3, CDC34, FBXO21, PSMD7, PSMD9, PSMD13, RPS27A, UBAP2L, UFD1, UFM1, USP10, and ZNF598; −7 to −40%, median = −20%).

#### Immune system, inflammatory response and apoptosis

Of the 538 regulated proteins between EW and LW, 26 were associated with inflammatory and immune function and 8 with apoptosis. More specifically, regulated proteins involved in the inflammatory process were very few (ITIH4, p=0.02; SELENOP, p=0.005; and FETUB, p=0.02) but were all 3 downregulated between EW and LW (respectively −33, −22 and −80%). Levels of 3 of the 8 proteins involved in apoptosis were lower in LW versus EW (−(24 to −52%; IER3IP1, p=0.03; PDCD5, p=0.02 and S100A14, p=0.001) while the other 5 were upregulated between EW and LW periods (+33, to +114%, median=+57%). Furthermore, 24 proteins involved in the immune system were significantly regulated in this dataset (12 proteins downregulated, −22 to −65%, median=−35%; 11 proteins upregulated, +21 to +119%, median=+66%; NCF1 not detected in EW). Interestingly, the complement cascade seemed to be clearly downregulated as 7 proteins involved in this process were significantly found in lower abundance in LW vs EW (−22 to −65%, median=−35%; CD59, p=0.005; C4A, p=0.03; C1QTNF9, p=0.04; C5, p=0.004; C8A, p=0.009; C3, p=0.03; C1RL, p=0.02). This pattern was confirmed at a less stringent significance threshold (0.05<p<0.1), where 5 additional proteins were downregulated while none of the proteins detected were upregulated (CFH, −40%, p=0.087; CFI, −42%, p=0.078; CFB, −49%, p=0.053; C9, −30%, p=0.075).

#### Carbohydrate metabolism and insulin signaling

Out of the 21 significantly regulated proteins classified in the carbohydrate metabolism pathway (Figures 3A and 3B), 14 were downregulated in LW vs EW (−15 to −56%, median=−37%), suggesting an overall reduction in carbohydrate metabolism in LW. More specifically, glucose uptake in hepatocytes did not seem to be altered as SLC2A2 expression was not changed between EW and LW (p=0.619). For glycolysis and gluconeogenesis processes, proteins such as GAPDH (−15%, p=0.024), GPI (−27%, p=0.021), MINPP1 (−30%, p=0.020), ALDOB (−37%, p=0.001), PFKP (−19%, p=0.044), PFKL (−31%, p=0.022) or FBP1 (−32% as an average of two isoforms, p=0.012 and p<0.0001) and were all showing lower levels in LW versus EW. Conversely, HK1 appeared upregulated in LW vs EW (+75%; p=0.008). Interestingly, animals exhibiting the highest expression levels of FBP1 corresponded to those showing the greatest fattening phenotype (positive correlation between BMC and FBP1 levels, p=0.01; Figure 3C). In addition, proteins involved in anaerobic glycolysis (LDHA: −51%, p=0.017 for one isoform), fructose biosynthesis (SORD, three isoforms: median=−51%, p=0.003 to p=0.017), xylose metabolism (XYLB: −18%, p=0.046), and glycerol degradation (GPD1: −38%, p=0.028; GPD2: −39%, p=0.013) were also downregulated between EW and LW, except for AQP9 which was upregulated (+29%, p=0.025). In contrast, proteins involved in glycogen metabolism, including PGM1 (two isoforms, +85% in average, p=0.019 and p=0.036) and PYGB (brain form; +31%, p=0.046) were upregulated in LW compared to EW (except for PYGL: liver form; −16%, p=0.026). A less stringent analysis (0.05<p<0.1) confirmed regulations of proteins involved in glycolysis and gluconeogenesis processes (ALDOA, +30%; p=0.052; GCK, +7%; p=0.089; and PGK1, +38%, p=0.098), glycerol degradation (GK, +38%, p=0.055), anaerobic glycolysis (second isoform of LDHA: −17%; p=0.08).

**Figure 3.**
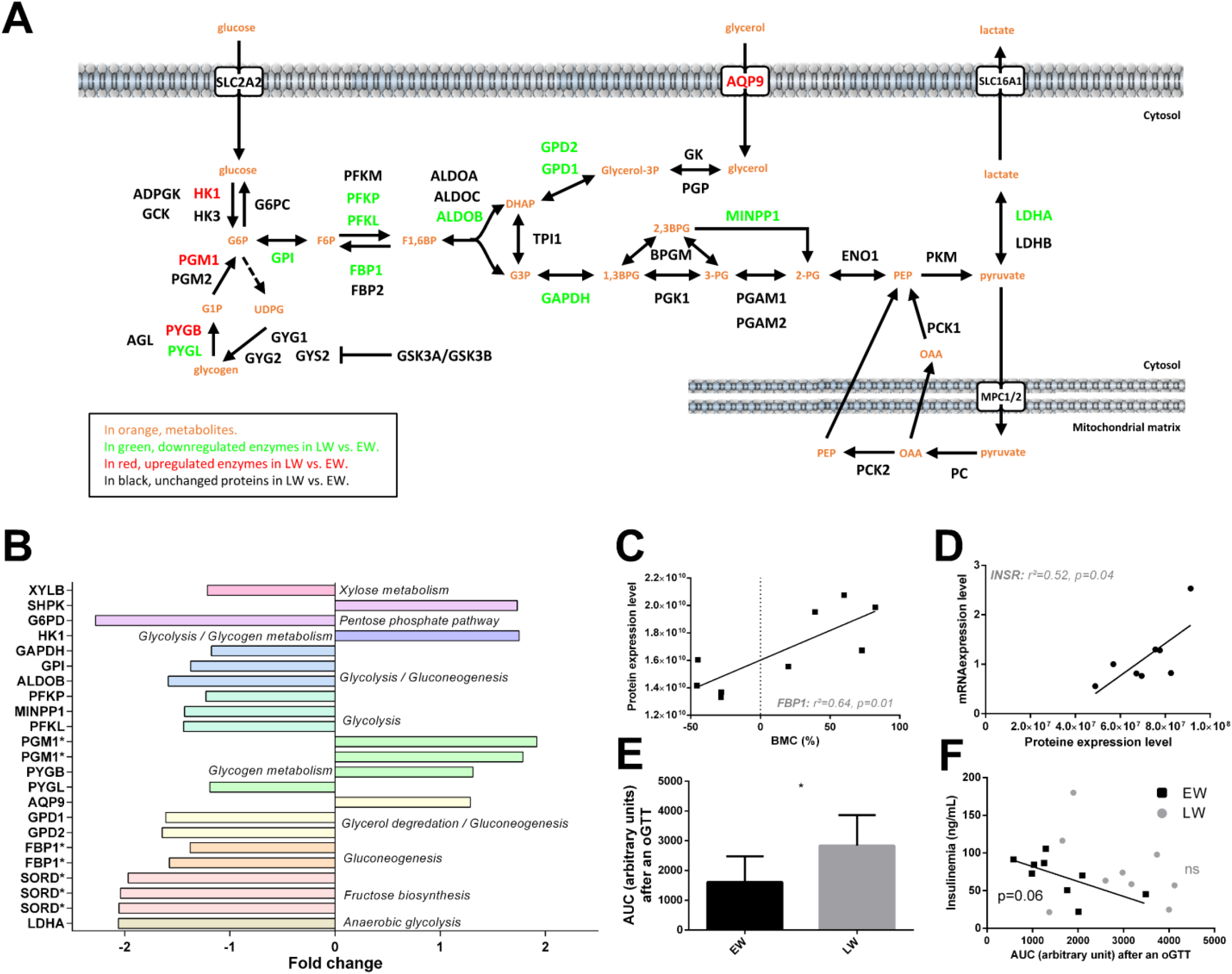
Regulation of liver carbohydrate metabolism and insulin signaling in grey mouse lemurs during winter. The relative abundance of liver proteins involved in carbohydrate metabolism measured in mouse lemurs (N=5/group) in early (EW) vs late (LW) winter is shown using a color code in the corresponding pathways (panel A), and mean fold changes in LW vs EW can be visualized in panel B (presence of several isoforms of the same protein are indicated with asterisks (*)). Significant and positive correlations were found between protein expression level of fructose-1,6-bisphosphatase 1 (FBP1) and body mass changes (BMC, panel C), and between mRNA and protein levels of insulin receptor (INSR, panel D). Retrieving data from a previous study showing the most extreme phenotypes (EW vs LW) revealed a greater intolerance to glucose in LW than in EW during an oral glucose tolerance tests (oGTT, panel E; results are presented as the means ± SEM), and a negative correlation between insulinemia and glucose intolerance (AUC, area under the curve) for EW lemurs, albeit not significant. * indicate significance at p < 0.05 between EW and LW.

In our dataset, no clear regulation of insulin signaling pathway was detected in the liver between EW and LW. Protein levels of INSR remained unchanged (+17%, p=0.86) although we observed a positive correlation between RNA and protein levels (p=0.04, Figure 3D). On another hand, IRS1 tended to be increased in LW animals (+30%, p=0.079), this upregulation being potentially sustained by reduced levels of MAPK9 (−27%, p=0.004), map kinases acting as repressors of IRS1 [30]. In accordance with the decreased utilization of glucose in LW suggested by the proteomic data, we observed a greater intolerance to glucose in LW than in EW animals resulting from an oral glucose tolerance test (oGTT) (Figure 3E). Interestingly, while circulating basal levels of insulin remained unchanged between EW and LW (data not shown), animals exhibiting the highest insulinemia during EW were the same animals showing the best regulation of glycaemia after oGTT (r^2^=0.42, p=0.057; Figure 3F); and this correlation was lost in LW (p=0.39; Figure 3F).

#### Lipid metabolism

Of the 47 significantly regulated proteins classified in the ‘lipid metabolism pathway’, the abundance of 24 of them was increased (+20 to +923%, median=+49%) and that of 23 others was decreased (−6 to −68%, median=−42%) during the EW period compared to the LW period (Figure 4A). Overall, liver lipid or fatty acid synthesis seemed to be slightly downregulated towards the end of winter (Figure 4A) as 55% of the proteins involved in these processes were decreased (e.g. HTD2, −28%, p=0.036; SCD, −81%, p=0.001; MCAT, −20%, p=0.003; HSD17B12, −33%, p=0.006; PHOSPHO1, −68%, p=0.008; HMGCS1, −66%, p=0.026; FDPS, −56%, p=0.006). Though, ACSL5 (+153%, p=0.033) and ACOT4 (+56%, p=0.034), which are specifically involved in the biosynthesis of unsaturated fatty acids (FA), were strongly upregulated in LW animals (Figure 4A). In this respect, the metabolism of arachidonic acid (C20:4 n-6) seemed to be altered during the winter period as CYP4F2 (+99%, p=0.019) and EPHX2 (+196%, p<0.0001) were highly and significantly upregulated in LW. No regulation was observed in alpha-linolenic acid metabolism pathway. Besides, apart from NSDHL (−61% in LW vs EW), proteins involved in liver CHOL (RALY, +29%, p=0.030) and bile acid (SLC27A5, +108%, p=0.026; CYP27A1, +38%, p=0.001; and HSD17B4, +21% on average for two isoforms, p=0.017 and p=0.029) synthesis were upregulated from the EW to LW. A less stringent analysis confirmed regulations of proteins involved in bile acid synthesis (ACOT8, −12%; BAAT, −22%; HSD3B7, +12%; ABCB11, +87%; SULT2A2, +205%).

**Figure 4.**
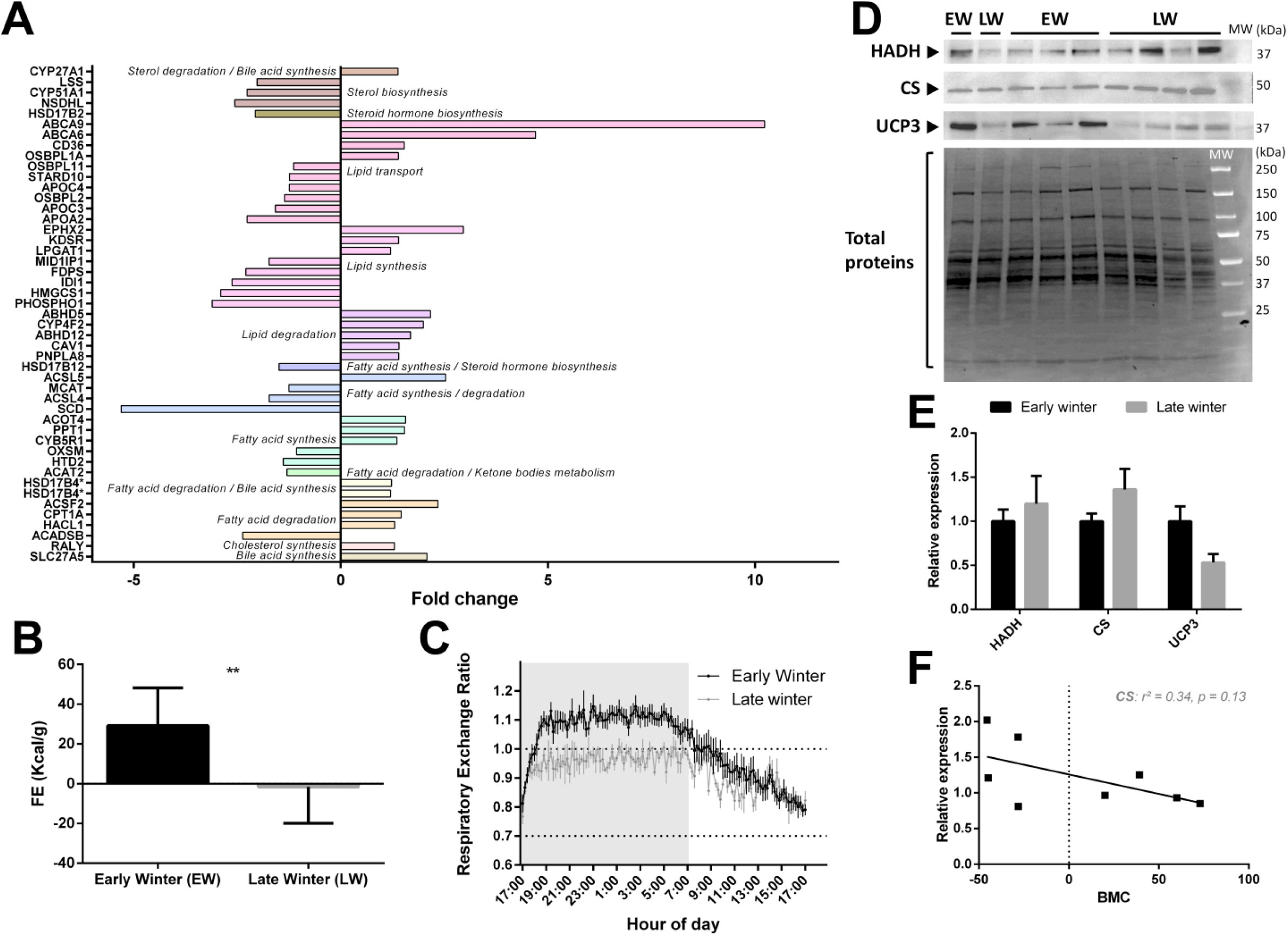
Regulation of liver lipid metabolism in grey mouse lemurs during winter. The relative abundance of liver proteins involved in carbohydrate metabolism in early (EW) and late (LW) winter mouse lemurs (N=5/group) is shown using mean fold changes in LW vs EW (panel A; presence of several isoforms of the same protein are indicated with asterisks (*). Food efficiency (FE), calculated as the ratio between food intake (in Kcal/day) and BMC, was significantly different between EW and LW animals (panel B), and the respiratory exchange ratio measured in EW (black line) vs LW (grey line) winter reflected the main energy substrates used depending on time-of-day (panel C). Protein expression levels of mitochondrial hydroxyacyl-coenzyme A dehydrogenase (HADH), citrate synthase (CS) and uncoupling protein 3 (UCP3) were measured using western-blot analysis in liver samples from mouse lemurs in EW and LW (blots shown in panel D and relative expression in panel E; N = 4-5/group; results are presented as the means ± SEM). Despite a trend toward a negative relationship, no significant correlation was obtained between protein levels of CS and body mass changes (BMC) (panel F). * and ** indicate significance at p < 0.05 and 0.01 between EW and LW, respectively.

In contrast, most of the proteins (77%; except ACADSB, −58%, p=0.003 and ACAT2, −23%, p=0.008) involved in FA or lipid degradation, including ABHD5 (+116%, p=0.037), ACSF2 (+134%, p=0.034), PNPLA8 (+40%, p=0.020), and CPT1A (+46%, p=0.040), were quantified as significantly increased in LW relative to EW. ACOX1, which is involved in the first step of beta-oxidation, was not significantly regulated at the protein level (p=0.348) but showed a strong negative correlation between mRNA and protein expression levels (r^2^=0.71, p=0.008). FA transport regulation was not regulated in a clear way as some proteins (4 out of 10) were upregulated (ABCA9, +923%, p=0.039; ABCA6, +370%, p=0.042; CD36, +53%, p=0.026; OSBPL1A, +39%, p=0.004) and some others (6 out of 10) were downregulated (e.g. APOA2, −56%, p=0.038; APOC3, −37%, p=0.035; OSBPL2, −27%, p=0.001; APOC4, −20%, p=0.042).

Observations at the proteome level accords well with variations observed at the physiological level in the individuals exhibiting the most extreme fattening (EW) and leaning (LW) phenotypes (see Methods). Indeed, animals in LW actually ate less than in EW (p=0.0003) and food efficiency was dramatically decreased to negative values (Figure 4B) and went along with a decrease of respiratory exchange ratio close to 0.7 (Figure 4C), therefore reflecting increased lipid mobilization from fat stores. Lipid oxidation in LW was not statistically supported by changes in liver HADH and CS (Figure 4D and 4E), although both proteins tended to be increased in LW vs EW (+20%, p=0.61 and +36%, p=0.23, respectively). However, animals showing the greatest body mass loss (corresponding to animals in LW) tended to be those exhibiting the greatest levels of CS (r^2^=0.34, p=0.13). Finally, mRNA levels of the transcription factor Xbp1 were not modified between EW and LW (Figure 4F) and the protein was not detected in the proteomic analysis.

#### Energy metabolism

Among the 538 significantly regulated proteins between EW and LW, 30 proteins were classified in the ‘Energy metabolism’ pathway, including 13 upregulated (+15 to +111%, median=+23%) and 17 downregulated proteins (−55 to −12%, median=−21%) in LW vs EW. A rapid overview of the energy metabolism pathway (Figure 5A) highlighted a possible downregulation of both tricarboxylic acid (TCA) cycle and mitochondrial respiration, in LW vs EW. The expression of 6 out of 8 proteins involved in TCA cycle (Figure 5B) was decreased in LW relative to EW. Indeed, in addition to ACLY (−62%, p=0.040) and SDHD (−18%, p=0.037), the expression levels of pyruvate dehydrogenases (PDHA1: −36%, p=0.001; PDHB: −36%, p=0.002; PDHX: −25%, p=0.002), as well as that of PDK1 (−54%, p<0.0001), which is known to inhibit pyruvate dehydrogenase and therefore to limit feeding of the TCA cycle with glycolytic intermediates, were lower in LW vs EW. By contrast, IDH2, which is involved in the interconversion of oxalosuccinate into either isocitrate or 2-oxo-glutarate, and SDHAF1, which is involved in the assembly of succinate dehydrogenase, were upregulated between EW and LW (+59%, p=0.011, and +10%, p=0.037, respectively).

**Figure 5.**
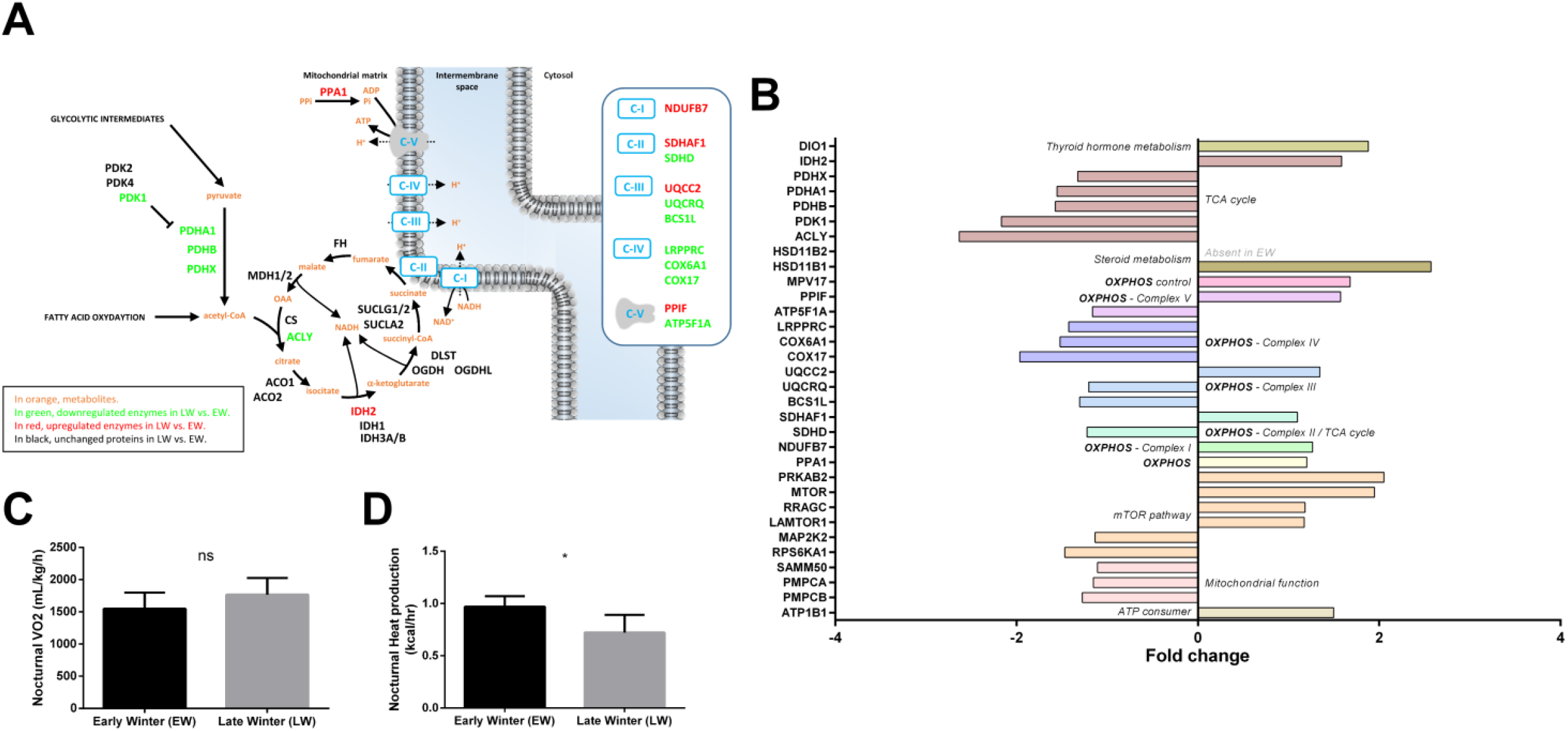
Regulation of liver energy metabolism in grey mouse lemurs during winter. The relative abundance of liver proteins involved in mitochondrial energy metabolism in early (EW) and late (LW) winter mouse lemurs (N=5/group) is shown using a color code in the corresponding pathways (panel A), and mean fold changes in LW vs EW can be visualized in panel B (HSD11B2 was not detected in EW). Although no change could be observed in nocturnal oxygen consumption (VO2) between EW and LW animals, nocturnal heat dissipation was significantly lower in LW lemurs (panel D). * indicate significance at p < 0.05 between EW and LW.

Proteins involved in mitochondrial oxidative phosphorylation (OXPHOS) showed a tendency to be downregulated in LW vs EW animals (Figure 5A and B, 7 out of 13 regulated proteins at p<0.05). In parallel, even if oxygen consumption remained unchanged between EW and LW (Figure 5C), heat production was significantly decreased in LW compared to EW (Figure 5D). Also, NDUFB7, a subunit of OXPHOS complex I, was significantly upregulated in EW animals (+27%, p=0.024). As already mentioned, complex II did not seem neither upregulated nor downregulated between EW and LW as SDHD decreased (−18%) while SDHAF1 increased (+10%). Two proteins constitutive of cytochrome c reductase (Complex III: UQCRQ, −17%, p=0.013) and cytochrome c oxidase (Complex IV: COX6A1, −34%, p=0.010) showed significantly decreased levels in LW vs EW. In addition, complex III assembly factors were either significantly downregulated (Complex III: BCS1L: −23%, p=0.036; Complex IV: COX17, −49%, p=0.022) or significantly upregulated (Complex III: UQCC2, +35%, p=0.003). Finally, one catalytic subunit of ATP synthase (Complex V) was significantly downregulated in LW vs EW (ATP5F1A, −14%, p=0.028), whereas the complex V assembly factor PPIF was upregulated (+57%, p=0.044). Interestingly, PPA1, a protein involved in fueling of ATP synthase with Pi (inorganic Phosphate) was significantly increased in LW vs EW animals (+20%, p=0.048). In addition, UCP3 was not detected in the proteomic analysis, but showed a significant downregulation in LW in the western blot analysis (−47%, p=0.04; Figure 4D and 4E). More generally, enzymes involved in mitochondrial processing peptidase (PMPCA: −13%, p=0.011; PMPCB: −21%, p=0.043) and SAMM50 (−10%, p=0.024) were less abundant in LW vs EW. Contrary to what was generally observed for mitochondrial proteins, we also could see that expression levels of important ATP consumers located at the plasma membrane were increase in LW (ATP1B1: +50%; p=0.038).

Metabolism is regulated by key factors like FOXO, MTOR and AMPK. While FOXO was not detected in our analysis, MTOR and AMPK signaling pathways seemed to be involved. Increased levels of MTOR itself was observed in LW (+95%; p=0.006). On the reverse the mTORC1 complex could be involved as suggested by the upregulation of the AMPK subunit PRKAB2 (+105%, p=0.002) and RRAGC (+18%, p=0.025). The increase RRAGC abundance in LW was paralleled by increased expression of LAMTOR1 (+17%, p=0.019), which is involved in amino acid sensing.

Finally, thyroid hormones and steroids are involved in key biological processes, including energy metabolism. While thyroid hormone signaling seemed to be increased in LW relative to EW animals (DIO1, +88%, p=0.013), steroid metabolism was overall enhanced at multiple levels. Indeed, HSD11B1 (+157%, p=0.038) and HSD11B2 (non-detected in EW), which are involved in cortisol synthesis and inactivation, were both significantly upregulated in LW animals. Less stringent statistical analysis confirmed significant regulations of TCA cycle (ACO1: −16%, p=0.052; MDH2: −17%, p=0.077; and IDH3B: −21%, p=0.079), OXPHOS (14 out of 21 proteins to be downregulated at p<0.1 including NDUFS5, NDUFA4, NDUFA2, SDHC, UQCRB, COX6B1 and COX7A2; Table S1) or ATP use (ATP1B3: +66%; p=0.081). In addition, the 30% increase in IRS1 levels (p=0.079), combined with the 18% decrease in PRKCA levels (p=0.089) in LW vs EW did not allow to determine if mTORC2 may play a role in mouse lemurs during winter. In contrast, regulations of RHEB (+10%, p=0.085) and RRAGA (+20%, p=0.076) tended to confirm a role of mTORC1 in the transition from EW to LW.

#### Amino acid metabolism

We observed changing levels for liver enzymes involved in the metabolism of several amino acids (Figure 6A). The liver is indeed important for directing the use of amino acids towards carbohydrate and/or lipid anabolic pathways. We notably observed a general lower abundance of many enzymes involved in pathways that make use of amino acids for glucose neosynthesis in LW vs EW animals (−21 to −60%; SDSL: −35%, p=0.024; AGXT, −23%, p=0.030; MAT1A, −24%, p=0.014; BHMT, −45%, p=0.025; AMDHD1, −41%, p=0.0009; HAL, −57%, p=0.001; CTH, −34% for one isoform, p=0.032; GLUD1, −28%, p=0.013; DDO, −63%, p=0.002; DMGDH, −46%, p=0.011). Only one enzyme of these latter pathways was higher abundant in LW (SHMT1, +30%, p=0.007). The enzymes involved in the synthesis of both carbohydrate and ketone bodies from amino acids manifested opposite regulations in LW (ACMSD, +122%, p=0.011GCDH, −30%, p=0.042; ACAT2: −23%, p=0.008). Several enzymes involved in ammonia detoxification and the urea cycle were found differentially expressed, notably GLUL (+197% in LW, p=0.038), and ASS1 (−24% in LW, p=0.015). Finally, the significant higher abundance of four other important proteins linked to amino acid metabolism was observed in LW (Figure 6A), including SMS (+50%, p=0.016), GRHPR (+26%, p=0.027), SAT2 (+41%, p=0.025) and DBT (+20%, p=0.0009); one additional protein was less abundant in LW (ALDH18A1, −33%, p=0.003).

**Figure 6.**
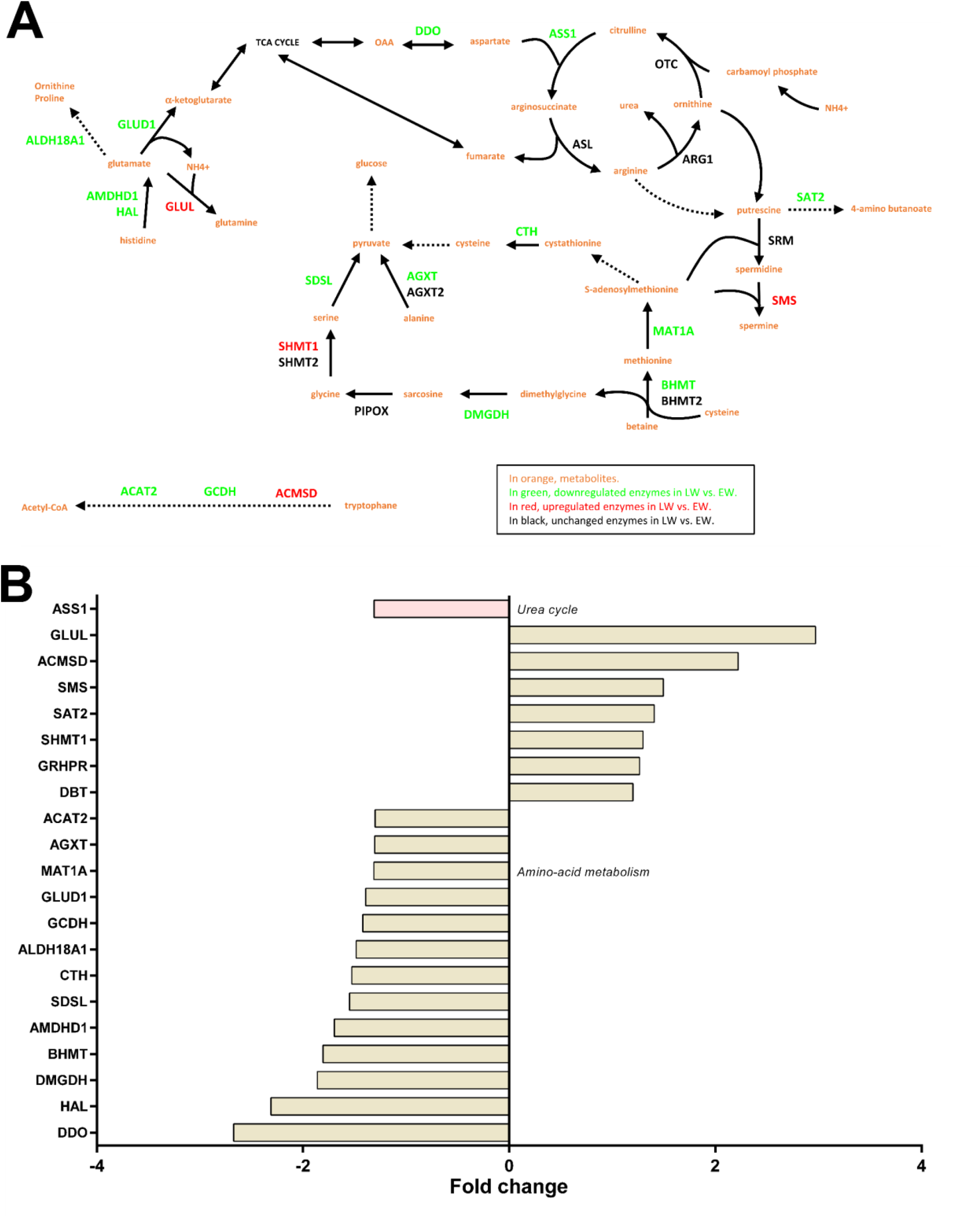
Regulation of liver amino acid metabolism and the urea cycle in grey mouse lemurs during winter. The relative abundance of liver proteins involved in amino acid metabolism and the urea cycle in early (EW) and late (LW) winter mouse lemurs (N=5/group) is shown using a color code in the corresponding pathways (panel A), and mean fold changes in LW vs EW can be visualized in panel B.

**Figure 7.**
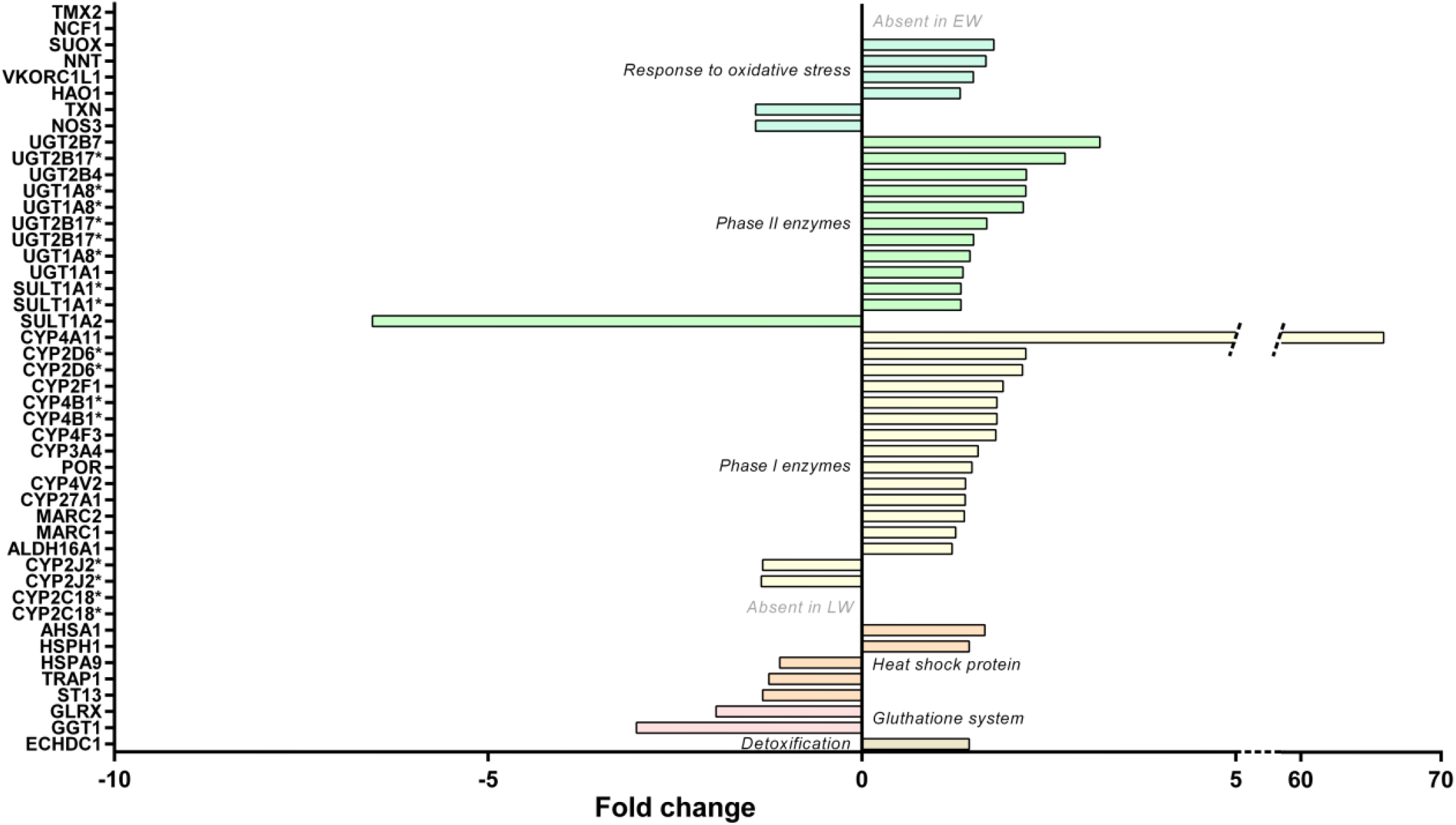
Regulation of liver stress responses in grey mouse lemurs during winter. The relative abundance of liver proteins involved in liver detoxification, oxidative stress, and that of heat shock proteins in early (EW) and late (LW) winter mouse lemurs (N=5/group) are shown using mean fold changes in LW vs EW. Presence of several isoforms of the same protein are indicated with asterisks (*).

Additional proteins were regulated in LW compared to EW when the p-value threshold was increased to p<0.1, notably several enzymes that regulate amino acid metabolism (CSAD, +28%, p=0.082; BCAT2, +9%, p=0.089; HPD, +29%, p=0.059; IVD, −45%, p=0.053), gluconeogenesis (PIPOX, +60%, p=0.073; and one isoform of CTH, −24%, p=0.085), and NAGS (−22% in LW, p=0.051).

#### Response to oxidative stress

The regulation of protein expressions involved in the response to free radicals did not clearly indicate changes in levels of liver oxidative stress between LW and EW (see Table S1). Indeed, proteins that favor free radical or hydrogen peroxide or nitric oxide production exhibited either higher (HAO1: +31%; p=0.031; and NCF1, only detected in LW) or lower (NOS3: −30%, p=0.037) expression levels in LW vs EW animals. Similarly, several antioxidant enzymes were found at higher expression levels in the liver during LW (VKORC1L1, +49%, p=0.032; and TMX2, uniquely detected in LW), whereas other proteins were determined as less expressed (TXN, −30%, p=0.030). Also, two proteins involved in glutathione system were less expressed in LW (GGT1, −67%, p=0.012; GLRX, −49%, p=0.028). Regarding heat shock proteins, which are oxidative stress-responsive proteins, higher levels during LW were recorded for two of them (AHSA1, +64%, p=0.001; HSPH1, +43%, p=0.048), and lower levels for three of them (ST13, −25%, p=0.012; TRAP1, −20%, p=0.007; HSPA9, −9%, p=0.038).

Additional changes were highlighted by using higher p-values (0.1<p<0.05), with higher expressed proteins in LW, including antioxidant enzymes (CAT, +24%, p=0.067; PRDX6, +31%, p=0.058; GCLC, +21%, p=0.055), and one heat shock protein (HSPA12A, +30%, p=0.071). In the same way, data were in line with a lower expression level of in LW for other antioxidant enzymes (HMOX2, −20%, p=0.087; PRDX4, −28%, p=0.052), as well as glutathione system factors (MGST1, −15%, p=0.066; GSTA5, −41%, p=0.088), and one chaperone protein (P4HB, −15%, p=0.087).

#### Liver detoxification

Examining the liver detoxification function, a majority of proteins exhibited higher abundances in LW vs EW (Figure 6B and Table S1), including 14 phase I enzymes (ALDH16A1, 2 isoforms of CYP2D6, CYP2F1, CYP27A1, CYP3A4, CYP4A11, 2 isoforms of CYP4B1, CYP4F3, CYP4V2, MARC1, MARC2, and POR; +21 to +6491%, median = +38%), 11 phase II enzymes (2 isoforms of SULT1A1, UGT1A1, 3 isoforms of UGT1A8, UGT2B4, UGT2B7, 3 isoforms of UGT2B17; +32 to +218%, median = +49%), and two other enzymes (ECHDC1 and SUOX, +31-77%, median = +43%). Nevertheless, other phase I enzymes or isoforms were quantified and exhibited downregulations in LW (2 isoforms of CYP2J2, −25 and −26%; and 2 isoforms of CYP2C18, not detected in LW – 6 other isoforms of CYP2C18 were also detected but not regulated, Table S1). This was also the case for an additional phase II enzyme (SULT1A2, −85%; p=0.008) with lower abundances in LW animals (Figure 6B). Again, being less stringent on the p-value threshold highlighted additional changes for phase I enzymes (CYP2C19, 2 upregulated isoforms in LW, +49% and +168%, and one downregulated isoform, −21%) and for one phase II enzyme (SULT2A1, +205% in LW).

## Discussion

Seasonal species are subject to very strong variations in environmental conditions, including energy availability. Amongst adaptive responses to such constraints, the energy-saving mechanisms, set up to survive periods of low energy availability, are coupled with preparatory phases during which organisms are storing a maximum of energy in the form of fat. These transitions from high to low food availability (and the opposite) trigger strong variations in body condition, which must be accompanied by profound molecular remodeling. Because the liver is acting as a main hub in metabolic regulation, we decided to investigate how the variations of the hepatic proteome relate with the phenotypic features of the mouse lemur between early winter (EW) and late winter (LW), i.e. fattening and leaning, respectively. Our results emphasize the regulations that operate in the liver between EW and LW in the mouse lemur, involving significant changes in glycolysis/gluconeogenesis, lipid metabolism but also detoxification. Although this study was conducted in captive animals, the regulations we identified reflected the seasonal phenotype of wild mouse lemurs and may emphasize protective mechanisms against metabolic disorders.

### Massive fattening does not induce a pathological state as witnessed by liver response

Seasonal species anticipate periods of low energy availability by storing large amounts of fat. Fattening results from the combination of food intake, though food availability begins to reduce in field conditions, and reduced metabolic rate. This process is obviously enhanced in captive conditions as animals do not experience reduction in food availability, and an increase in food intake has in fact been reported previously in EW vs. LW animals [11]. Considering the known causes for hypertriglyceridemia [31], the elevated circulating levels of triglycerides we observed in captive animals during EW (Figure 1C) are probably due to a higher food intake. From body mass dynamics and food efficiency values, fattening is obvious in EW, and the activity of lipoprotein lipase is likely expected to be high at this stage for favoring triglyceride hydrolysis from chylomicrons and very low-density lipoproteins (VLDL) and their delivery to white adipose tissue. The production of chylomicrons might therefore exceed triglyceride clearance in EW animals. It is worth noting here that our macroscopic observation of the livers (not shown) suggests that this situation of enhanced lipid accretion does not become pathogenic as no ectopic accumulation of fat was observed in the liver of EW lemurs, even with a diet that was rich in carbohydrates. This result is in line with the absence of any increase in hepatic lipid levels in rats fed a high-carb diet [32]. Conversely, hepatic lipogenesis is induced in healthy subjects in response to a high carbohydrate diet, which involves an increase in protein expression levels of transcription factor XBP1 [33]. In mouse lemurs, hepatic Xbp1 mRNA expression (Figure 4F) remained at similar levels in EW and LW, suggesting a lack of response to food intake levels (higher in EW) and a regulation uncoupled from the global orientation of lipid metabolism (lipogenesis in EW versus lipolysis and lipid oxidation in LW). Our proteomic data in EW versus LW, which highlight higher abundances of several hepatic proteins involved in lipid and fatty acid synthesis as well as lipid transport, does not necessarily contradict this hypothesis. Although the regulations we observe are supporting the hypothesis that the physiology of EW animals is mainly oriented towards the constitution of fat reserves, it is unlikely that this actually reflects an increased production of VLDL in EW. Indeed, the major component of VLDL, namely apolipoprotein B-100 [34], remained at stable levels in our two groups of lemurs (see Table S1). In addition, an increased production of VLDL would have resulted in the increased generation of LDL particles [35], and our data show the opposite. LDL particles are the main drivers for atherogenic disorders [36], therefore, their lower levels in EW animals are another supporting element that fattening process in EW animals is not leading to a pathogenic situation. Hence, proteome regulations for lipid metabolism should rather be considered in the light of the inhibition of lipogenesis in LW which should contribute to promoting the mobilization and oxidation of lipids (see below).

In human adipose tissue, de novo lipogenesis is activated after high carbohydrate feeding [37], and our results support that similar regulations happen in EW animals. The obvious conversion of the excess of ingested carbs into fat reserves in captive EW mouse lemurs is testified by RER values above 1 during the night (feeding phase) [38,39]. In the liver of EW animals, the lower abundances of enzymes involved in the synthesis of unsaturated fatty acids (ACSL5 and ACOT4) and of enzymes known to convert arachidonic acid (C20:4 n-6) (CYP4F2 and EPHX2) suggest a reduced turnover for this fatty acid and a possible maintenance of its concentration at high levels. This would be consistent with a previous study in female mouse lemurs in EW, where n-6 fatty acids represented ~39% of total lipids in the liver (vs ~13% for n-3), with C20:4 n-6 being the main form [26]. The maintenance of high content of n-6 fatty acids (and reduced content of n-3 fatty acids) was notably associated with deep and long torpor bouts, and it is also observed in hibernating species [40–42]. Therefore, high levels of n-6 fatty acids in EW mouse lemurs could support their extensive use of daily torpor. Conversely the higher levels of CYP4F2 in LW versus EW animals may serve other functions (see below).

Interestingly, it has be demonstrated that captive mouse lemurs reduce their oxygen consumption during the first half of winter [11], however, we show here that this global reduction in metabolic rate does not translate into a reduction in nocturnal oxygen consumption or in liver protein levels that play a prominent role in mitochondrial processes for energy fuel oxidation(TCA cycle and OXPHOS). These last results corroborate with a study in golden-mantled ground squirrels [43], which highlighted that respiratory rates (all respiratory states) reached their highest levels during pre-hibernation in fattening animals. Expression of proteins involved in OXPHOS activity (Complex I and Complex I+II) was greatest during this period, suggesting a higher hepatic carbohydrate oxidation capacity, which is in line with our results (see below). Hence, the global decrease in oxygen consumption in EW animals might be mostly attributed to drastic reduction of muscle activity and muscle metabolism instead of liver regulations. Our results, consisting in higher levels of glycolytic factors and lower levels of fatty acid oxidation-related proteins, suggest that maintenance of liver metabolic activity in EW animals and the maintained production of liver ATP largely involves carbohydrate fuels. This hypothesis is in perfect line with the preservation of insulin sensitivity during the fattening phase as it has been reported previously [11] and it is further confirmed in the current study based on oGTT results and the well-regulated glycaemia. Previous studies showed an induction of insulin resistance during high-carbohydrate diets that involved not only hepatic steatosis but also the induction of *G6pc* and *Gckr* genes as a protective mechanism promoting homeostasis [44]. In our data, no such changes were observed in EW animals concerning liver lipid accumulation (see above) and regarding protein expression levels of G6PC and GCKR (see Table S1). EW mouse lemurs therefore exhibit a physiological state that appears to be safely managed without the induction of the known adverse effects that are usually triggered by high carbohydrate feeding or in relation with fat accumulation in white adipose. Also, at the systemic level, massive fattening did not induce excess of fat deposits in metabolic tissues (absence of hepatic steatosis; personal observation), and gene [11] and protein expression (present study) did not evidence any sign of hepatic inflammation. In that sense, EW animals look very similar to animals in the pre-hibernating period.

### Liver metabolism in LW is sustained by mobilization of peripheral lipid stores, but independent of glucose metabolism

The massive fattening observed in male mouse lemurs in the first weeks of winter naturally stops in the middle of winter. Afterwards, the combination of decreased food intake (despite no change in food availability) and increased locomotor activity and metabolic rates [11] results in a decrease in body mass. Such seasonal regulation of feeding behavior and satiety signals has already been demonstrated in hibernating species and seems to be controlled through modifications in hormonal signaling [45] and/or the gut-brain axis [46]. The reduction in RER in LW (Figure 4C) testifies that oxidative metabolism in LW animals is essentially fueled from the mobilization of fat stores instead of food carbs. Though, levels of RER do not reach a strict value of 0.7, meaning that metabolic oxidation is not purely lipidic, which is probably linked to the fact that animals are still eating high-carbs food (food intake is not null). The higher levels of CYP4F2 in LW animals may promote conversion of C20:4 n-6 into 20-HETE, thus allowing for adjustment of liver metabolic activity to control fat-dependent energy supply and metabolism [47]. The action of CYP4F2 may also favor the catabolism of pro-inflammatory eicosanoids, such as leukotriene B4 [48], which could prevent the development of an inflammation due to large lipid fluxes in LW mouse lemurs as already suggested [11]. Similar regulations have already been described in hibernating animals, albeit with slight differences between animal species [49]. In another hand, it is possible that liver n-6 fatty acids in LW are used in relation with the reactivation of the reproductive axis in male mouse lemurs at the end of winter (see below). Indeed, C20:4 n-6 is a precursor for prostaglandins that play a role in reproduction and in thermoregulation [50,51]. At the proteomic level, the switch from carbs to lipid mobilization in LW might involve a reduction in glucose utilization in the liver, as suggested by a decreased abundance of glycolytic enzymes and of LDHA. In the same way, reduced expression levels of several isoforms of pyruvate dehydrogenase (PDHA1, PDHB, and PDHX) suggest a reduction of the oxidation of glycolytic intermediates. Conversely, beta-oxidation of fatty acids is nicely reflected by an increased abundance (although not significant) of HADH and citrate synthase (CS) in LW. These protein regulations are in perfect line with observations at the systemic level in LW mouse lemurs, i.e., a lower tolerance to glucose after an oGTT albeit basal insulin levels are higher than in EW animals [11]. On the other hand, the reduced levels of fructose-1,6-bisphosphatase 1 (FBP1) would be in line with a reduction in liver gluconeogenesis in LW compared to EW.

Glycerol is one of the main sources for liver gluconeogenesis, after entry into hepatocytes through aquaporin-9 (AQP9) known as a key rate-limited step [52]. In LW animals, glycerol release due to white adipose tissue lipolysis and its uptake in liver cells are expected to increase. Our data showing higher expression levels of AQP9 protein during LW support this hypothesis. However, the reduced abundances of GPD1 and GPD2 suggest that glycerol oxidation might not be increased in the liver of LW animals. Because metabolism is oriented toward fat utilization at this stage, it is unlikely that glycerol is used for liver lipogenesis and, therefore, glycerol might accumulate in LW. Strikingly, increased expression of AQP9 at the mRNA level has been linked to insulin deficiency or to insulin resistance in mice [53]. It is therefore possible that the regulation of AQP9 in LW lemurs may be linked to glucose intolerance development (as reflected by oGTT results) and linked to a possible alteration in insulin signaling [11]. However, insulinemia and insulin receptor (INSR) expression levels remain unchanged between EW and LW in the current study, which suggests that glucose intolerance may not be due to an alteration in liver insulin signaling or that the cause of such an alteration (if it actually happens) are to be searched elsewhere (see below).

Amino acids are another main substrate for liver gluconeogenesis. An increase in amino acid-derived hepatic gluconeogenesis was well described in response to fasting, as well as the resulting enhancement of protein protection through the maintenance of IGF-1 levels [54]. Such maintenance of IGF-1 abundance was also observed during caloric restriction in mouse lemurs [55] and let us hypothesize that this process is involved in the transition between EW and LW hepatic phenotypes. However, the reduced expression of all enzymes involved in liver gluconeogenesis from amino acids, between EW and LW, is another evidence that liver glucose metabolism is not favored in LW. An association has been made between increased fasting-induced autophagy and increased hepatic gluconeogenesis [56], that contributes to maintain a basal level of glycaemia. However, such mechanisms do not appear to be involved in maintaining the circulating glucose levels that have been observed in mouse lemurs between EW and LW [11]. Indeed, the molecular regulations we observed point toward a reduction in gluconeogenesis in LW (see above) but contradictory data were recorded in the current study for autophagy process. In fact, key autophagy factors (RMC1, BNIP3, and SNAP29) were more abundant in the liver of LW versus EW animals, while at the same time expression levels of WD repeat-containing protein 6 (WDR6), a negative regulator of amino acid starvation-induced autophagy (McKnight et al., 2012), were increased between EW and LW (Table S1; p=0.003). In accordance with the apparent repression of liver glucose metabolism in LW, glucose intolerance, associated with higher basal levels of insulin, was observed in LW in this species [11]. That suggests a possible impairment in insulin sensitivity during fat mobilization in LW. Furthermore, the expression of FBP1 protein is lower at the end of winter, when the animal is leaning, compared to the beginning of winter. According to the literature, FBP1 is involved in the negative regulation of appetite and of adiposity [57], and it prevents the development of insulin resistance [58]. Indeed, increased expression of FBP1 in liver after nutrient excess increases circulating satiety hormones and reduces appetite-stimulating neuropeptides and thus seems to provide a feedback mechanism to limit weight gain. In LW, animals are probably more starved than in EW, which could explain the reduced expression of FBP1, and further explain the transient settlement of insulin resistance in LW. However, pathway analysis from our proteomic dataset did not show any major regulation in the insulin signaling pathway, although the vast majority of actors were well detected. We think that repression of liver glucose metabolism in LW, without an important alteration of liver insulin sensitivity might indicate, again, that a pathological stage is not reached in this tissue. It would be interesting to pursue our investigation by focusing on proteome regulations in skeletal muscle to better identify seasonal regulation of insulin signaling. In addition, seasonal control of this function could come from finest regulations, including phosphorylation of key proteins [59].

Despite a decrease in blood urea nitrogen during LW [11], it is difficult to conclude if the liver urea cycle is significantly affected in LW mouse lemurs. Indeed, we observed an upregulation of two key enzymes (DDO and ASS1) whereas expression of three other enzymes (OTC, ARG1 and ASL) remained unchanged between EW and LW. As explained above, we cannot exclude a finest regulation led by post-translational modifications and/or modulation of the uptake of amino acids may be involved in downregulation of the cycle, which should be explored in future studies. A possible hypothesis is that the rate of flux of amino acids toward the liver, a key regulator of ureagenesis, may likely be low in LW due to a decrease in the muscle efflux of amino acids. This idea is supported by the self-imposed ‘caloric restriction’ situation (according to the fact that food intake spontaneously decreases) in LW, which seems to resemble the low energy diets in several animals, including food-storing hibernators, which maintain their protein pool and limit muscle loss [10]. Hence, the reduction in food intake, together with increased energy expenditure (see below), is accompanied with the mobilization of fat stores in LW lemurs, which might help spare body proteins. Further, increased locomotor activity in LW animals (data not shown) might even favor physical exercise-stimulating protein synthesis [60]. Future studies to characterize the molecular events in the muscle from LW versus EW lemurs, with emphasis on the regulation of protein balance, should help answer this question.

Finally, although an increased oxygen consumption was observed in LW [11], the regulations at the protein level observed here for the different subunits of the mitochondrial respiratory chain complexes suggest a general decrease in liver OXPHOS activity. At the systemic level, the increase in VO_2_ in LW could be explained by increased locomotor activity [20], and by a higher muscle metabolic rate. We expect that muscle proteome regulations would reveal changes in OXPHOS activity, and enhanced burning of fat to fuel metabolic processes. On another hand, hepatic protein regulations from our data set suggest that mitochondrial uncoupling may be decreased in LW (UCP3 being downregulated as compared to EW), which conforms to the increase in lipid beta-oxidation (CPT1 upregulation in LW).

In this study, we identified a hepatic proteome downregulation of liver gluconeogenesis in LW that is not fully in line with regulations observed during hibernation, e.g. in hibernating bears [61]. In addition, species arousing from hibernation exhibit an increase in hepatic uncoupling [62], which was not observed in LW mouse lemurs. It is crucial to distinct mouse lemurs from these other species knowing that they are still eating during LW and may not fully rely on their fat reserves. Such phenotype may however rather resemble that of food-storing hibernators, which rely on food reserves during hibernation, therefore restoring blood glucose levels through food intake during arousals [10].

### Fat mobilization in LW is associated with reactivation of the reproductive system and liver detoxification

In addition to the metabolic phenotype observed in wintering male mouse lemurs, the activity of the reproductive system is also deeply modified. Indeed, while a complete regression of testes and a decrease in circulating levels of testosterone are observed in EW, testicular recrudescence begins ~20 weeks after the beginning of winter time and is sustained by an increase in testosterone blood levels [20,63,64]. Body mass loss in LW is probably mostly explained by increased metabolic rates that could be a consequence of a more important locomotor activity [11]. However, we suspect that reactivation of the reproductive system (which can be likened to a yearly puberty) may contribute to body mass regulations and has been neglected so far. The net energetic cost of spermatogenesis is largely overlooked in mammals, while few studies showed that this process may explain body mass variations in reptiles [65]. In addition, male grey mouse lemurs present all the features describing sperm competition: high spermatogenesis efficacy and motility, and low percentage of defect [66,67]. Most of the reproductive constraint in the male mouse lemur rely on the access to fertile females, at the very beginning of the summer season. These observations may highlight an anticipatory phenomenon of sexual recrudescence in LW that would result in a significant energy expenditure. This also means that there is a strong need for producing testosterone in LW, whereas EW animals experience complete damping in circulating testosterone levels. Interestingly, the hepatic regulation of proteins involved in CHOL and steroid metabolism may reflect this process. Indeed, excess circulating CHOL in LW does not seem to be paralleled by an increase in its delivery to the liver by HDL. The augmented levels of circulating LDL could positively regulate CHOL delivery to the gonads for testosterone production but it can also reflect a situation at risk for the cardiovascular system of mouse lemurs during LW. However, the absence of significant variation in the ratio CHOL/HDL between EW and LW (although close to significance), and the maintenance of this ratio at a low level throughout winter [68], leads to conclude that the proportion of so-called « bad cholesterol » (LDL) might remain stable and low. The fact that liver low-density lipoprotein receptor (LDLR) and hepatic triacylglycerol lipase (LIPC) protein abundances remain unchanged between EW and LW does not support an increase in liver CHOL uptake. However, bile acid synthesis and secretion could be increased in LW animals, since we observed a significant upregulation for several key proteins of this pathway (CYP27A1, CYP3A4, HSD17B4, SLC27A5, and UGT2B4). It may be that increased bile acid production in LW may help remove excess CHOL, and it may also constitute an answer to the reduction of food intake for improving dietary lipid absorption, a well-known function of bile [69]. Interestingly, an activation of the bile acid receptor (NR1H4 or FXR) has been reported to repress gluconeogenic gene expressions and to enhance insulin sensitivity in diabetic mice [70]. In the same way, the possible overproduction of bile acids in LW lemurs may contribute to the global downregulation of liver gluconeogenesis we observed (see above). Alternatively, liver interconversion of cholesterol into bile acids has been linked to increased adaptive thermogenesis involving brown-adipose tissue (BAT) [71]. An increased activity in this thermogenic tissue has been observed in LW in mouse lemurs [11] and may explain the possible higher synthesis of bile acids in LW.

One last hepatic-specific process that seemed notably regulated in LW in mouse lemur is detoxification. With regards to oxidative stress, although systemic DNA damage (measuring urinary 8-OHdG levels) and circulating gamma-glutamyl transpeptidase (aka GGT) have both been demonstrated as increased in LW in mouse lemurs [11], proteome liver regulations of main antioxidant enzymes were unchanged or even reduced. Further evidence suggests that systemic DNA damage is related to metabolic rates in male mouse lemurs [72] and could not necessarily translate into a hepatic signature. Such observation was already made in the arctic ground squirrel, where oxidative stress during hibernation is tissue-specific, being observed in BAT (a thermogenic tissue) but not in liver [73]. The absence of liver oxidative stress upregulation in LW could indicate an absence of pathogenic state for mouse lemurs. It is worth noting here that this may be a peculiar feature in mouse lemurs as it was already pointed that prolonged fasting in rats triggers an increase in liver oxidative stress [74]. This appears further reinforced by the apparent correct functioning of liver protein folding processes, as suggested by the fact that very few proteins involved in protein processing in endoplasmic reticulum were regulated in LW vs EW animals, except from downregulation of MAN1B1 and protein OS-9 that function in quality control targeting of misfolded proteins for degradation.

In our study, we successfully quantified 27 out of 30 detected proteins known to be involved in detoxification and all of them, mostly phase I and II enzymes, were upregulated between EW and LW. This may reflect an important activation of this process in LW. Aldehyde dehydrogenases are proteins helping to detoxify the accumulated derivatives of lipid peroxidation during torpor [75], and a previous study already revealed their upregulation in golden-mantled ground squirrels at the entrance in hibernation [76] and after the fattening phase [77]. Interestingly, golden-mantled ground squirrels are described as food-storing hibernators, they mostly rely on their fat reserves during hibernation [78], therefore resembling mouse lemurs’ phenotype in LW [11]. On the other hand, studies in hibernating bears suggest a down-regulation of liver proteins involved in detoxification due to their prolonged fasted state [79]. Based on these observations, we hypothesized that enhancement of liver detoxification in LW mouse lemurs could highlight their greater sensitivity to xenobiotics as compared to the EW situation. In fact, all the animals used for the proteomics analysis were anesthetized using a 2% isoflurane / 98% air mixture for ~45 minutes (after ketamine induction). The impact of a general gaseous anesthesia was already demonstrated on hepatic gene expressions of drug metabolizing enzymes in rats after long-lasting procedure (6h under anesthesia) [80], although this has not yet been investigated for shorter exposure. . According to the striking physiological differences that exist between EW and LW, which are supported by hepatic proteome regulations we identified, we cannot reject the hypothesis that sensitivity to isoflurane per se may differ during winter and be greater in LW vs EW. Alternatively, enhanced sensitivity to xenobiotics in LW might correspond to an anticipating process to the upcoming summer time with higher food intake. Anticipation processes are important during nutritional transitions. For instance, intestinal villi atrophy induced by food deprivation on the short term is restored during late fasting in laboratory rats, through cell divisions and a temporary interruption of apoptosis that correspond to anticipation of refeeding and the need for improving food assimilation and the restoration of the whole body condition [81].

## Conclusion

During winter, grey mouse lemurs (*Microcebus murinus*) exhibit an early phase of fattening followed by a late phase of fat mobilization. By investigating how their liver proteome is modulated from EW to LW, we could show that hepatic regulations nicely support fuel partitioning at the molecular level, and really reflect the metabolic phenotype. Moreover, no sign of the typical adverse effects that are associated with fattening in humans, e.g. high levels of LDL or the absence of signs of altered insulin signaling and inflammation in the liver, were observed in EW animals. Similarly, despite glucose intolerance as indicated from an oGTT, we did not identify evidence of the alteration of insulin signaling in LW at the hepatic level. Animal species living in seasonal environments with strong energetic constraints have developed unique adaptive mechanisms conferring resistance capabilities to safely face metabolic challenges. Hence, this work perfectly illustrates that studying such species is relevant to identify targets involved in metabolic flexibility and energy homeostasis, and could be an important asset to the biomedical field.

## Acknowledgment

We are grateful to Dr. P. Guterl for his help in bioinformatics analysis of proteomics data.

## Funding

This work was financially supported by the CNRS (2014 PEPS ExoMod program) and Strasbourg University (H2E project; IdEx Unistra) and the French Proteomic Infrastructure (ProFI; ANR-10-INSB-08-03). During the tenure of this study, M.B.D. was the recipient of a PhD grant from the University of Strasbourg (IdEx Unistra), and C.B. from the French Ministry of Higher Education, Research and Innovation.

## Competing interests

We declare no competing interests.

## Author contributions

J.T. conceived the study and carried out the experimental work on mouse lemurs. F.B. supervised proteomics analyses. B.C. and M.B.D. conducted mass spectrometry-based analyses, and CB western-blot analyses. J.T. and F.B. drafted the manuscript. All authors gave final approval for publication.

## Data Availability

The mass spectrometry proteomics data have been deposited to the ProteomeXchange Consortium via the PRIDE partner repository with the dataset identifier PXD030198. Electronic supplementary materials are provided for the detailed mass spectrometry-based identification and quantification of all proteins (Table S1) and the principal component analysis (PCA) of protein abundances (Table S2). Mouse lemur individual data remain available upon demand to the corresponding authors.

## Ethics approval

All experiments were carried out in accordance with the European Communities Council Directive (86/609/EEC) and all experimental procedures were evaluated by an independent ethical council and approved for ethical contentment by the French government (authorizations n°03210.02, n°APAFIS#3004-2015111015031850 and n°APAFIS#3697-2016012111304236).

## Notes

### Competing Interest Statement

The authors have declared no competing interest.

